# Microtubule-based Transport is Essential to Distribute RNA and Nascent Protein in Skeletal Muscle

**DOI:** 10.1101/2021.02.26.433059

**Authors:** Lance T. Denes, Chase P. Kelley, Eric T. Wang

## Abstract

While the importance of RNA localization in highly differentiated cells is well appreciated, basic principles of RNA localization in skeletal muscle remain poorly characterized. Here, we develop a method to detect single RNA molecules and quantify localization patterns in skeletal myofibers, and we uncover a critical and general role for directed transport of RNPs in muscle. We find that RNAs are localized and translated along cytoskeletal filaments, and we identify the Z-disk as a biological hub for RNA localization and protein synthesis. We show that muscle development triggers complete reliance on the lattice-like microtubule network to transport RNAs and that disruption of microtubules leads to striking accumulation of RNPs and nascent protein around myonuclei. Our observations suggest that active transport may be globally required to distribute RNAs in highly differentiated cells and reveal fundamental mechanisms relevant to myopathies caused by perturbations to RNPs, microtubules, and the nuclear envelope.

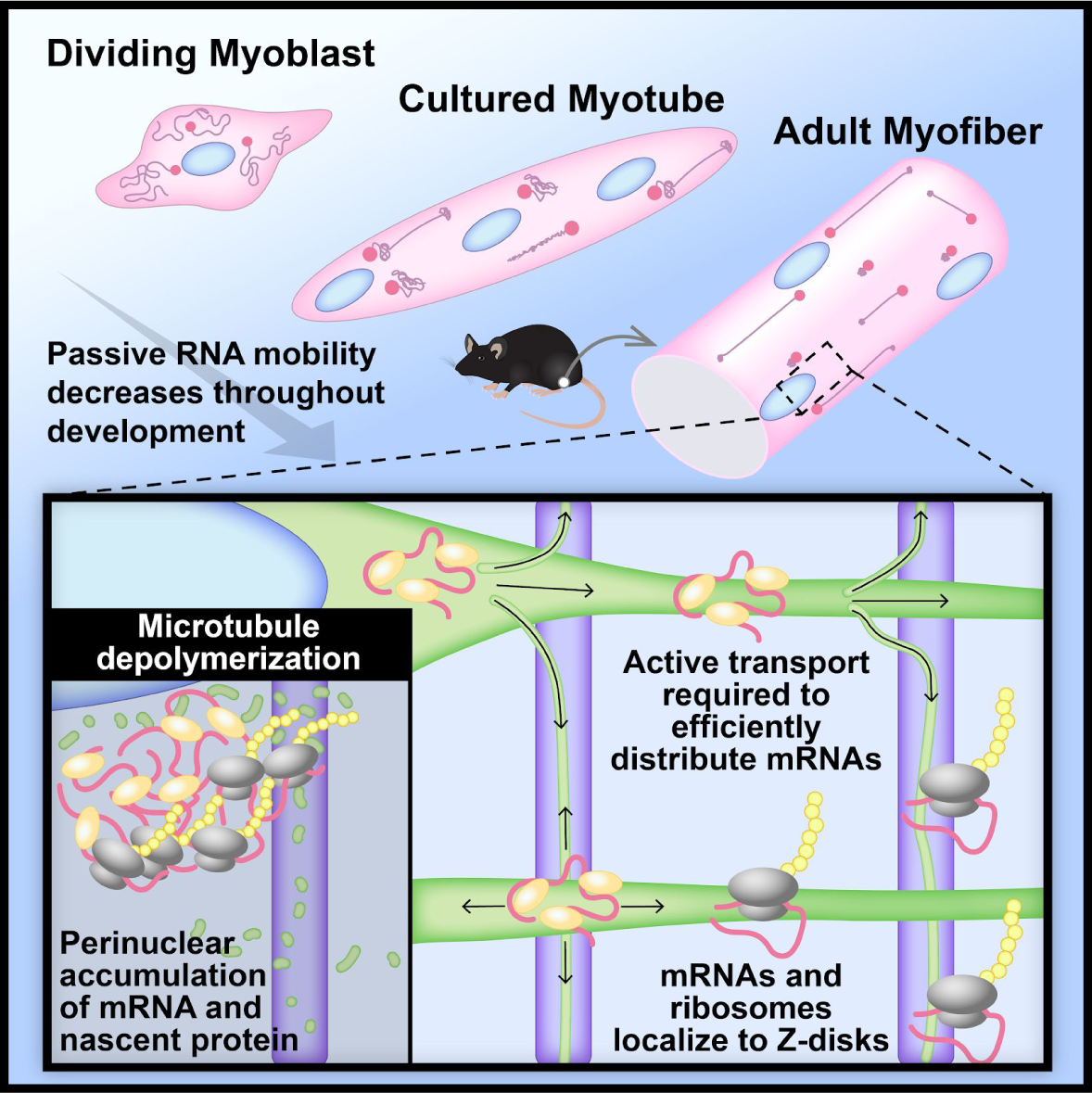

## INTRODUCTION

The spatial distribution of RNA influences compartmentalized protein synthesis, responsiveness to local stimuli, and cell polarity (Martin and Ephrussi, 2009). Proper RNA localization patterns are required for mating type specification in yeast (Long et al., 1997), *Drosophila* embryo patterning (Lécuyer et al., 2007), and memory consolidation in the mammalian brain, among other functions (Dalla Costa et al., 2021; Miller et al., 2002). RNA *cis*-elements recruit specific RNA-binding proteins (RBPs) to form higher-order ribonucleoprotein (RNP) granules, together dictating their biophysical properties and interacting partners (Weber and Brangwynne, 2012). Mechanisms for localizing RNPs include diffusion and entrapment (Forrest and Gavis, 2003), local stabilization (Bashirullah et al., 2001), and active transport (Park et al., 2014). To ensure local production of proteins, RNAs are often translationally repressed while in transit but subsequently unmasked by spatiotemporal cues at their final destination (Graber et al., 2013; Kiebler and Bassell, 2006). RNA localization has been especially well-studied in neurons, where local synthesis of proteins at synapses is required for proper function (Dalla Costa et al., 2021). Directed transport is essential to rapidly deliver RNAs to thousands of neuronal synapses, sometimes as far as a meter away from the nucleus (Holt et al., 2019). To put this in perspective, it has been estimated that a β-actin mRNA would take 48 days to arrive at the tip of a 500 µm-long axon by diffusion alone (Turner-Bridger et al., 2018).

Similar to neurons, striated skeletal muscle cells, or myofibers, are large, post-mitotic, and highly differentiated (Bentzinger et al., 2012)—but in contrast to neurons, they are syncytial tubes containing hundreds of myonuclei (Bruusgaard et al., 2006). In healthy adult muscle, myonuclei are typically located at the periphery of the myofiber and are evenly spaced apart; this configuration is proposed to maximize efficiency of gene expression (Bruusgaard et al., 2003; Hansson et al., 2020). Myofibers also contain clustered myonuclei (Bruusgaard et al., 2003) at myotendinous junctions (MTJs), where myofibers interface with tendons (Charvet et al., 2012), and at the neuromuscular junction (NMJ), where motor neurons innervate (Wu et al., 2010). Myonuclear density varies between muscle types: in mouse *extensor digitorum longus*, nuclei are ∼30 µm apart, and the volume of syncytial cytoplasm per nucleus, often referred to as the “myonuclear domain”, is ∼5x the volume of a fibroblast (Hansson et al., 2020; Swanson et al., 1991). Similar to neurons, myofibers must populate a large cytoplasmic space with gene products, but also face the added challenge of coordinating multiple nuclei within a shared cytoplasm. While this problem has been appreciated for decades (Pavlath et al., 1989), the mechanisms by which RNAs from each nucleus distribute within that space are poorly understood. Enrichment of specific RNAs at MTJs (Dix and Eisenberg, 1990) and NMJs (Sanes et al., 1991) has been reported; in the case of the NMJ, these patterns are in part driven by unique postsynaptic transcriptional programs (Hippenmeyer et al., 2007). In the extra-synaptic region, various RNA localization patterns have been described, some of which appear contradictory: 1) confinement near nuclei (Nevalainen et al., 2013; Ralston and Hall, 1992; Ralston et al., 1997), 2) enrichment at the myofiber surface (Dix and Eisenberg, 1988; Mitsui et al., 1997; Nissinen et al., 2005), 3) uniform dispersion between nuclei and within the myofiber core (Kann and Krauss, 2019; Shoemaker et al., 1999), and 4) association with cytoskeletal filaments or striated patterning (Cripe et al., 1993; Fulton and Alftine, 1997; Pomeroy et al., 1991). Notably, many of these studies were performed in cultured cells and/or employed low resolution autoradiographic techniques, and modern single molecule fluorescent *in situ* hybridization (smFISH) techniques have not yet been widely applied to adult myofibers, in part due to high levels of background autofluorescence.

A substantial portion of gene expression activity in myofibers supports the capacity to generate contractile force. The basic contractile unit in striated muscle cells, including skeletal myofibers and cardiac muscle cells (cardiomyocytes), is the sarcomere—an intricate, ∼2-µm-long cytoskeletal apparatus (Henderson et al., 2017). The sarcomere is anchored by Z-disks on either end and contains actin thin filaments and myosin thick filaments that slide over each other during contraction (Luther, 2009). The cytoplasm of a muscle cell is packed with sarcomeres, which are bundled into myofibrils, creating a dense yet highly ordered environment that restricts diffusion of large macromolecules (Papadopoulos et al., 2000). The importance of the sarcomere for muscle function is highlighted by frequent involvement of sarcomere proteins in skeletal and cardiac muscle disease (Kamisago et al., 2000; Sewry et al., 2019). A link between sarcomere assembly and RNA localization was proposed decades ago (Isaacs and Fulton, 1987), and has recently been supported by observations of protein synthesis machinery and mRNAs encoding sarcomere proteins at Z-disks in cardiomyocytes (Lewis et al., 2018; Rudolph et al., 2019).

In neurons, RNPs are actively transported along microtubules by the motor proteins kinesin and dynein, and these interactions are mediated by protein-based (Baumann et al., 2020; Dictenberg et al., 2008; Wu et al., 2020) or vesicular adaptors (Cioni et al., 2019; Liao et al., 2019). Properly regulated microtubule-based RNP transport and local translation are important to prevent neurological disease, which is often caused by mutations to microtubule associated proteins (Deniston et al., 2020; Dubey et al., 2015), motor proteins and adaptors (Hirokawa et al., 2010; Liao et al., 2019; Nicolas et al., 2018; Zhao et al., 2001), RBPs (Fernandopulle et al., 2021; Ramaswami et al., 2013), and regulators of translation (Banerjee et al., 2018; Kapur et al., 2017). In myofibers, the microtubule network forms a grid-like lattice that is interwoven throughout sarcomeres. Large bundles of antiparallel microtubule filaments extend from longitudinal poles of myonuclei, and smaller perpendicular bundles branch off along sarcomere Z-disks (Fig. S1A and Video S1) (Oddoux et al., 2013). The skeletal muscle microtubule network plays important roles in nuclear positioning (Cadot et al., 2012; Metzger et al., 2012), vesicular transport (McDade and Michele, 2014), and mechanotransduction (Kerr et al., 2015). Microtubules also play critical roles in similar processes in cardiomyocytes; accordingly, microtubule perturbations are implicated in both skeletal and cardiac myopathies (Caporizzo et al., 2019; Khairallah et al., 2012). Interestingly, localization of RNAs to the periphery of cardiomyocytes requires microtubules (Perhonen et al., 1998; Scholz et al., 2006), suggesting a role for active transport in moving RNAs through the cytoplasm of striated muscle cells. Cardiomyocytes are smaller than myofibers and typically contain only one or two nuclei (Landim-Vieira et al., 2020); thus, the role of active RNA transport in large and highly multinucleated skeletal myofibers remains unclear.

Similarly to neurons, aberrant RNA metabolism underlies a variety of striated muscle diseases (Banerjee et al., 2013; Guo et al., 2012; Kanadia et al., 2003; Picchiarelli and Dupuis, 2020; Zarnescu and Gregorio, 2013), and some RBPs, including TDP-43 and MBNL, are involved in disease pathogenesis in both muscle and neurons (Thornton, 2014; Weihl et al., 2008). Despite a potential role for RNPs to regulate the unique morphological and functional demands of muscle, few investigations have been conducted on RNA localization in myofibers. As a result, we have limited understanding of principles of RNP transport in this tissue and whether they parallel those observed in other cell types.

Here, we develop methods to image single RNA molecules together with protein markers in *ex vivo* skeletal myofibers, and we characterize localization patterns of a diverse set of RNAs. We uncover an intimate relationship between RNAs, sarcomeres, and the microtubule network, and we show that muscle development triggers an absolute reliance on microtubules to disperse mRNAs and avoid accumulation of large RNP granules near myonuclei. We discover that protein synthesis also occurs along cytoskeletal networks, but that efficient RNA transport does not depend on translation. By observing motile RNP granules in live myotubes, we identify diffusive and directed transport states, estimate motion parameters, and confirm by computational simulation that observed RNA localization patterns in myofibers can only be achieved by a significant directed transport component. These observations outline principles of RNA localization in muscle, with broad implications for RNA-protein homeostasis and overall muscle function.

## RESULTS

### RNAs Are Dispersed Throughout Skeletal Myofibers

We first developed a platform for robust quantitative analysis of subcellular RNA localization patterns in skeletal muscle. We studied *extensor digitorum longus* myofibers isolated from adult FVB/NJ mice (Fig. 1A) rather than tissue sections to avoid cryosectioning artifacts. Additionally, myofibers can be cultured for days and subjected to pharmacological treatments (Pasut et al., 2013). We initially attempted smiFISH to detect single mRNA molecules (Tsanov et al., 2016), but we found that at least 90 primary oligonucleotide probes per gene were required to overcome the strong autofluorescence in myofibers, making it intractable to visualize most mRNAs. We therefore implemented *in situ* hybridization chain reaction (HCR v3.0) technology (Choi et al., 2018) to amplify signal from a smaller number of primary probes, allowing us to visualize single mRNA molecules in myofibers with as few as 20 oligonucleotides (Fig. 1A). To quantitate localization patterns, we developed a robust computational pipeline that segmented myofibers and myonuclei from confocal image stacks and detected the positions of FISH spots (Fig. S1B). This pipeline allowed us to count the number of mRNAs in myofibers and measure their subcellular localization relative to structures in the myofiber (e.g. myonuclei or cytoskeleton).

**Figure 1.**
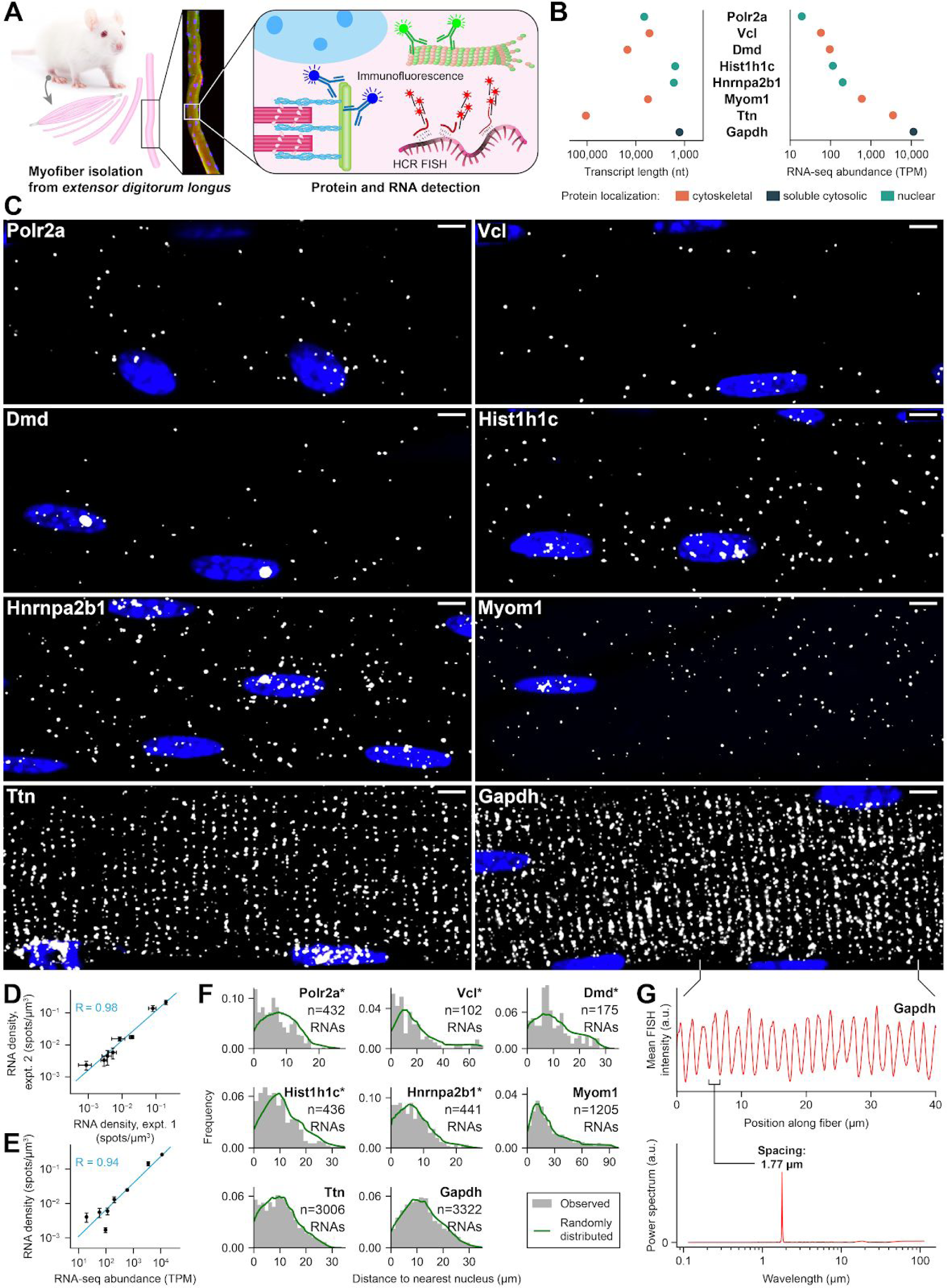
Single mRNAs Can Be Reproducibly and Accurately Detected by HCR smFISH and Are Dispersed Throughout Myofibers. A) Schematic describing experimental strategy to label RNAs and proteins of interest in adult skeletal muscle. Myofibers were isolated from adult mouse *extensor digitorum longus* and fixed immediately or cultured. RNAs of interest were labeled by smFISH using *in situ* hybridization chain reaction (HCR v3.0) technology for signal amplification. Proteins of interest were labeled by immunofluorescence (IF) (see Methods). See also Fig. S1A, Video S1, and Video S2. B) Transcript length (nucleotides, nt) and abundance in *tibialis anterior* muscle (transcripts per million, TPM) for mRNAs from each gene studied; colors represent encoded protein localization. C) FISH of mRNAs from each gene studied in isolated myofibers. Nuclei are labeled with DAPI. Scale bars: 5 μm. D) For each gene (black point), cytoplasmic mRNA density (spots/µm^3^) measured from FISH images using our computational pipeline was compared across separate experiments. Mean ± s.d. of at least n=3 images of individual myofibers per gene per experiment. Trendline: LLS regression; pearson R=0.98; p<0.05, Wald test. See also Fig. S1B. E) Cytoplasmic mRNA density was compared with transcripts per million (TPM) values from a *tibialis anterior* RNAseq dataset. Mean ± s.d of at least n=3 images of individual myofibers per gene. Trendline: LLS regression; pearson R=0.94; p<0.05, Wald test. F) Distance from cytoplasmic mRNAs to nearest nucleus (grey bars) measured for each gene studied and compared to a null distribution of randomly selected cytoplasmic coordinates (green lines). Distributions derived from images of at least two FISH-labeled myofibers per gene. Seven of eight genes were closer to nuclei than by chance, but the shift in median distance was never >3 µm. *p<0.05 with shift in median distance >0.5 µm, Mann-Whitney *U* test. See also Fig. S1C and S1D. G) Mean Gapdh FISH signal intensity of myofiber in (C) plotted for 40 µm along the longitudinal axis (top). The power spectral density of this signal calculated by discrete fast Fourier transform (bottom).

Using this platform, we studied a diverse set of eight genes to determine whether gene-specific properties such as gene expression level, transcript length, or encoded protein function were related to mRNA localization patterns (Fig. 1B). This set included mRNAs encoding sarcomere proteins (Ttn, Myom1), costamere proteins (Vcl, Dmd), a nuclear/cytoplasmic shuttling RNA-binding protein (Hnrnpa2b1), a histone protein (Hist1h1c), a subunit of RNA polymerase II (Polr2a), and a metabolic protein (Gapdh). All studied mRNAs are 5’-capped and polyadenylated, including Hist1h1c, which is polyadenylated in differentiated cells (Lyons et al., 2016).

We observed discrete FISH spots throughout the cytoplasm and nuclei of myofibers for all genes studied (Fig. 1C and Video S2). FISH spot densities (spots/μm^3^) were highly reproducible across experiments (Fig. 1D) and strongly correlated with mRNA copy numbers estimated from RNA-seq (Fig. 1E). Strikingly, we found that cytoplasmic mRNAs were not confined within individual myonuclear domains, but appeared well dispersed between myonuclei, indicating that mRNAs frequently travel at least tens of microns from progenitor nuclei. The number and size of intranuclear spots was variable between genes and may reflect differences in pre-mRNA transcription or processing rates for each gene. Large “blobs” were observed in some nuclei and were especially prominent for Ttn and Dmd mRNA; similar blobs have been detected at sites of transcription in pre-mRNA-specific FISH (Femino et al., 1998). We did not confine our probes to 5’ or 3’ ends of mRNAs; thus, they likely detected both pre-mRNAs lingering at sites of transcription as well as mature mRNAs still present in the nucleus. The large size of intranuclear Ttn and Dmd blobs may reflect long residence time of pre-mRNAs at transcription sites for these exceptionally large genes. Finally, the observation of only one large Dmd blob in each nucleus is consistent with a single site of transcription on the X chromosome.

To quantify the dispersion of each mRNA throughout the cytoplasm, we measured the distance from each cytoplasmic FISH spot to the nearest nucleus and compared these distances to a null distribution generated from random cytoplasmic coordinates. We found that RNAs from all genes except Myom1 were closer to nuclei on average than expected by chance (p < 0.05 by Mann-Whitney *U* test), but the effect was subtle, and RNAs from all genes populated regions of the myofiber cytoplasm furthest from nuclei (Fig. 1F). Specifically, the difference in median distance between experimental and randomized distributions in all cases was <3 μm, or ∼10% of the mean internuclear distance (Bruusgaard et al., 2003), and below 0.5 μm for Gapdh and Ttn. The magnitude of this shift was inversely correlated with RNA copy number (Pearson R = −0.79), uncorrelated with RNA length (Pearson R = −0.11), and not strictly related to encoded protein function (Fig. S1C). Additionally, we detected all RNAs at the myofiber surface and in the core, including Dmd and Vcl, which encode surface-localized proteins (Fig. S1D). Overall, our observations suggest that, regardless of function and localization of the encoded proteins, RNAs are well dispersed throughout the myofiber syncytium.

### RNAs Associate With Z-Disks and Microtubules

Ttn mRNA in the cytoplasm showed a clear striated pattern, consistent with previous observations in cultured myotubes (Fulton and Alftine, 1997) and cardiomyocytes (Rudolph et al., 2019), but we were surprised to find that Gapdh mRNA was similarly striated. We quantified the periodicity of the FISH signal for Gapdh by Fourier analysis, and we found that the peak wavelength of this pattern corresponded to the length of the sarcomere (Fig. 1G). The striated patterns for Gapdh and Ttn mRNAs motivated us to directly assess whether these and other mRNAs might lie along Z-disks, M-lines, or regions in between. We developed an immunofluorescence (IF)/HCR smFISH protocol to co-label sarcomeric proteins and RNAs of interest. We found that a strategy implementing IF first, followed by an additional fixation step before FISH, yielded robust detection of mRNAs and proteins for all combinations tested (see Methods). Because all RNAs studied could travel tens of microns away from myonuclei, and microtubules are involved in RNA transport in other cell types, we also investigated whether mRNAs might associate with microtubules by co-staining tubulin. IF labeling of telethonin and tubulin revealed robust Z-disks and an impressive microtubule lattice, respectively (Fig. S1A and Video S1). Consistent with previous studies, we observed large microtubule bundles extending from poles of myonuclei along the longitudinal axis, as well as smaller bundles branching out perpendicularly along the sarcomere Z-disk.

Surprisingly, we found that RNAs from *all* genes studied were clearly associated with both Z-disks and microtubules (Fig. 2A and 2B). In the absence of sarcomere and microtubule labeling, this localization pattern was visually apparent for highly expressed genes, but co-labeling of protein markers and mRNAs revealed this association for every gene studied, regardless of expression level. To quantitate the extent of association, we generated binary segmentations from microtubule and Z-disk image stacks and applied our computational pipeline to measure the distance from each cytoplasmic FISH spot to the nearest Z-disk and microtubule. We compared these distance distributions to a null distribution derived by repeatedly placing the same number of spots in random positions within the cytoplasm, and we found that mRNAs from all genes were significantly associated with Z-disks and microtubules (p < 0.05 by Mann-Whitney *U* test, Fig. 2D). We also noticed a preferential localization of mRNAs near perpendicular Z-disk-microtubule intersections (ZMIs). We categorized spots as “cytoskeleton-associated” if they were within 2 pixels (∼0.1 μm) of Z-disks or microtubules, and we measured the distance from each of these spots to the nearest ZMI; all cytoskeleton-associated mRNAs were slightly and significantly biased towards ZMIs as compared to a set of spots randomly placed along the cytoskeleton (p < 0.05 by Mann-Whitney *U* test, Fig. 2E). Overall, we were surprised to find that Z-disk localization was not limited to mRNAs encoding sarcomere proteins, but also included mRNAs encoding nuclear proteins. Even Myom1 mRNA, which encodes the M-line-localized protein myomesin-1, was localized to the Z-disk. Thus, localization of mRNAs to Z-disks appears to be a general rule in skeletal myofibers and not exclusively related to sarcomere assembly. In addition, association of RNAs with microtubules and ZMIs suggests they may be actively transported from nuclei to Z-disks along microtubules.

**Figure 2.**
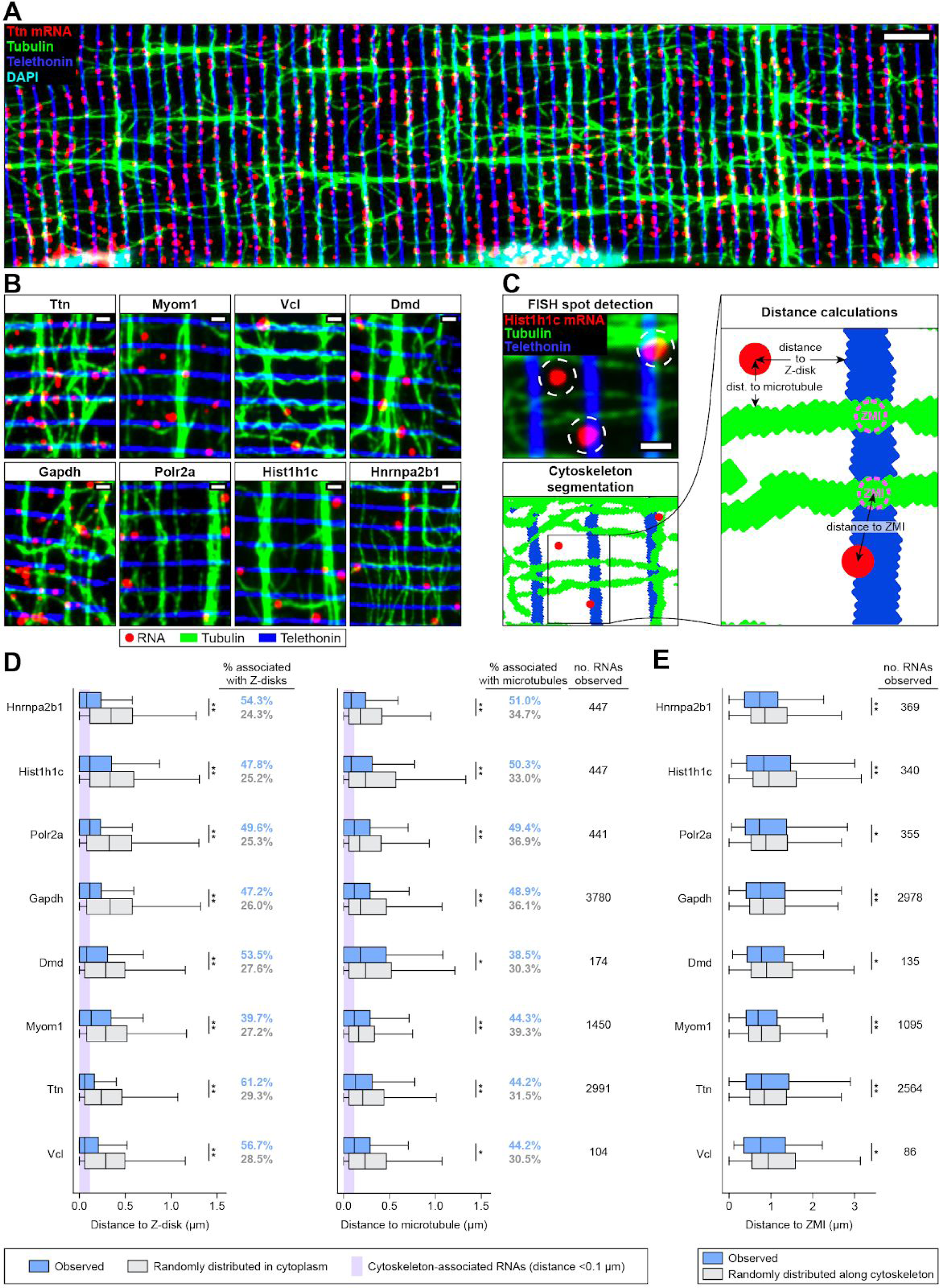
RNAs Associate With Z-disks and Microtubules. A) IF/FISH co-labeling of Ttn mRNAs (red), tubulin protein (microtubules, green) and telethonin protein (Z-disks, blue) in an isolated myofiber. Scale bar: 5 µm. B) Zoomed-in regions of IF/FISH co-labeled myofibers as in (A) for each gene studied. Scale bars: 1 µm. See also Fig. S2A. C) Schematic describing the computational pipeline used to assess mRNA-cytoskeleton association from images of IF/FISH co-labeled myofibers (A). mRNA transcript coordinates were identified, Z-disks and microtubules were segmented to generate binary masks, and distances from each mRNA molecule to the nearest Z-disk and microtubule were measured. Z-disk-microtubule intersections (ZMIs) were computationally detected, and the distance from each cytoskeleton-associated RNA to the nearest ZMI was measured. D) mRNAs from all eight genes studied were significantly associated with Z-disks and microtubules. Distribution of distances from cytoplasmic mRNAs (blue boxes) to Z-disks (left) and microtubules (right) compared to null distributions generated from randomly selected cytoplasmic coordinates (grey boxes). Data were derived from images of at least two FISH-labeled myofibers per gene for each gene studied. mRNAs located within 2 pixels (∼0.1 μm) of either filament were considered “cytoskeleton-associated” (purple shaded region, percentages). *p < 0.03. **p < 10^-4^, Mann-Whitney *U* test. E) Cytoskeleton-associated mRNAs from all genes studied were significantly biased towards ZMIs. Distribution of distances from cytoskeleton-associated mRNAs to ZMIs (blue boxes) compared to a null distribution generated from randomly selected coordinates along the cytoskeleton (grey boxes) for each of the eight genes studied. *p < 0.03, **p < 10^-4^, Mann-Whitney *U* test.

### NMJ-Specific RNAs Associate with Z-disks and Microtubules Around the NMJ

The eight mRNAs studied above are transcribed in myonuclei throughout the entire myofiber syncytium. At the NMJ, a cluster of specialized myonuclei at the postsynaptic membrane transcribe a distinct set of mRNAs under control of NMJ-specific transcription factors (Hippenmeyer et al., 2007). We asked how the localization of NMJ-expressed Chrne mRNA compared to the mRNAs we studied in the extra-synaptic myofiber. As expected, Chrne mRNA was highly concentrated in the immediate vicinity of the NMJ (Fig. S3A). Interestingly, it was also present in adjacent regions of the myofiber, including within extra-synaptic nuclei neighboring the NMJ, potentially indicating extra-synaptic sites of transcription. We imaged the NMJ region at 100x magnification from two different angles to obtain axial and radial cross-sections (Fig. S3B and S3C). While Chrne mRNA was highly enriched near the postsynaptic membrane, we also detected RNAs in the myofiber core that were preferentially associated with Z-disks and microtubules, similar to all other genes previously studied (Fig. S3B). We also found that, in comparison to extra-junctional regions, post-synaptic microtubules did not exhibit the grid-like organization observed in the extra-synaptic myofiber and instead formed a denser, nest-like matrix encasing the cluster of myonuclei (Fig. S3C). Chrne mRNAs were frequently localized within these dense bundles of microtubules. These data suggest that the nest-like NMJ microtubule network may restrict the mobility of RNAs produced in post-synaptic nuclei, although some may escape into the extra-synaptic cytoplasm and associate with Z-disks and microtubules there, similar to other extra-synaptic RNAs. Evidence of Chrne transcription in extra-synaptic nuclei also suggests that post-synaptic transcription factors (either the proteins or the RNAs that encode them) that drive Chrne expression may travel out of the postsynaptic region to nearby extra-synaptic nuclei.

### Distribution of mRNAs Throughout the Cytoplasm is Completely Dependent on Microtubules

Our observations show that RNAs are dispersed tens of microns away from myonuclei and also co-localize with microtubules and Z-disks in the cytoplasm. Furthermore, RNAs transcribed in post-synaptic nuclei are locally confined in a dense, nest-like microtubule network, but can also disperse and localize along the microtubule lattice outside this region. Thus, we hypothesized that microtubules may play a direct role in distributing RNAs from myonuclei. To investigate this possibility, we cultured myofibers in the presence of nocodazole, an antineoplastic agent that potently and reversibly depolymerizes microtubules and notably does not affect the sarcomere in myofibers (Fig. 3A and 3B) (Oddoux et al., 2013). Strikingly, after culturing myofibers in 5 µg/mL nocodazole for 18 hr, we found that mRNAs from all genes studied had strongly accumulated around the nuclear periphery. We then allowed microtubules to re-polymerize by washing out the nocodazole and culturing in fresh medium for an additional 4 hr. After 4 hr of washout, accumulated mRNAs were cleared from the nuclear periphery (Fig. 3C, 3D, and S4A).

**Figure 3.**
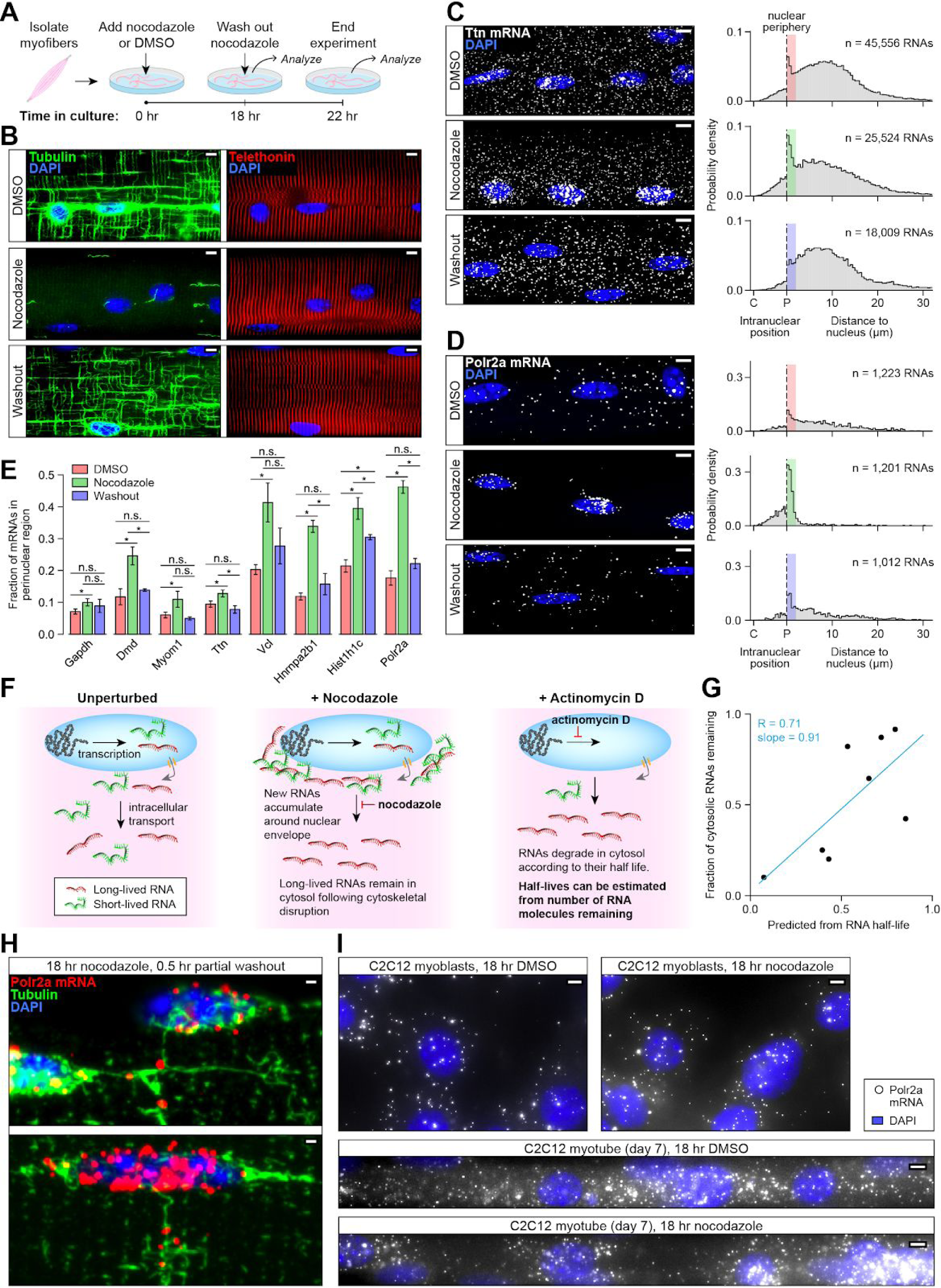
Distribution of RNAs Throughout the Cytoplasm Requires Microtubules. A) Schematic describing microtubule depolymerization experiment. Live myofibers were isolated and cultured with 5 μg/mL nocodazole or DMSO (control) for 18 hr. At 18 hr, one set of myofibers was collected, fixed, and processed for FISH to detect mRNAs from each gene. In another set of myofibers, nocodazole was washed out, allowing microtubules to re-polymerize, and fibers were cultured for an additional 4 hr before fixation and FISH. B) IF of tubulin protein (microtubules, green) and telethonin protein (Z-disks, red) in myofibers at each stage of the experiment described in (A). Microtubules depolymerize following nocodazole treatment and re-polymerize following nocodazole washout. Z-disks are not affected. Scale bars: 5 µm. C) FISH of Ttn mRNA in myofibers from the experimental time course described in (A) (left). Intranuclear position relative to centroid (C) and periphery (P) measured for mRNAs in the nucleus, and distance to nearest nucleus measured for mRNAs in the cytoplasm (right). Data derived from multiple images of FISH-labeled myofibers in each of two independent experiments. Perinuclear region (defined as <2 μm from the nuclear periphery) analyzed in (E) denoted by bars colored by experimental condition. Scale bars: 5 µm. D) Same as (C) but Polr2a mRNA. Scale bars: 5 µm. E) Fraction of mRNAs in the perinuclear region increased after 18 hr nocodazole treatment and was reduced after washout for all eight genes studied. Quantitation of the fraction of total mRNAs in the perinuclear region (colored bars in C, D, S4B) for each gene studied from the experimental time course described in (A). Mean ± SEM. Between n=2 and n=12 images of individual myofibers from two independent experiments were used for each gene/condition combination. *p < 0.05, Mann-Whitney *U* test. See also Fig. S4A. F) Schematic of the method using actinomycin D used to investigate cytosolic mRNA transcripts that persist following nocodazole treatment. For each gene studied, mRNA half-life was estimated from FISH spot densities measured after 0, 6, and 22 hr of transcriptional inhibition with actinomycin D. Half-lives were used to predict the fraction of cytoplasmic transcripts that should remain after 18 hr of nocodazole treatment if new transcripts were unable to exit the nucleus. See also Fig. S5A and S5B. G) For each of the eight genes studied, the fraction of mRNAs remaining in the cytoplasm after 18 hr nocodazole treatment (“Fraction cytosolic mRNAs remaining”, Y-axis) was correlated with the fraction predicted by the decay model (“Predicted from RNA half-life”, X-axis) outlined in (F). Trend line: LLS regression, slope=0.9; pearson R=0.71; p=0.045, Wald test. H) IF/FISH of tubulin protein (microtubules, green) and Polr2a mRNA (red) in myofibers cultured in nocodazole for 18 hr, followed by partial washout and 30 min of additional culture. Polr2a mRNAs were observed in the cytoplasm only at the ends of individual nucleus-connected microtubule segments. Scale bars: 1 µm. I) FISH of Polr2a mRNA in C2C12 myoblasts treated with DMSO or nocodazole for 18 hr, and in C2C12 myotubes differentiated for 7 days on micropatterned gelatin, then treated with DMSO or nocodazole for 18 hr. Scale bars: 5 µm.

We used our computational pipeline to rigorously quantitate the effect of nocodazole on mRNA localization. For cytoplasmic FISH spots, we measured their distance to the nearest nucleus, and for nuclear spots, we measured their intranuclear position relative to the centroid and periphery of the nucleus (Fig. 3C, 3D and S4A). We found that the fraction of mRNAs in the perinuclear region (within 2 μm of the nuclear periphery) was increased for all genes in nocodazole-treated myofibers relative to control (all p < 0.05 except Myom1, with p = 0.055 by Mann-Whitney *U* test, Fig. 3E). After 4 hr of nocodazole washout, the difference in fraction of perinuclear mRNAs relative to baseline was no longer detectable for any gene (p > 0.05, Mann-Whitney *U* test) except Hist1h1c, but Hist1h1c did show a significant decrease between nocodazole treatment and washout (p < 0.05, Mann-Whitney *U* test). We also observed some accumulation of RNAs at the intranuclear periphery, which was cleared after microtubule re-polymerization (Fig. 3C, 3D and S4A). Overall, the effect of microtubule depolymerisation was extreme and demonstrates a complete inability for exported RNAs to passively diffuse away from nuclei.

While RNAs from all eight genes accumulated in the perinuclear region, the fraction of RNAs depleted from the cytoplasm after nocodazole treatment varied between genes (Fig. 3C, 3D, and S4A). We hypothesized that these differences reflect differential rates of mRNA decay among genes, rather than differences in the ability of new mRNAs to enter the cytoplasm without microtubules (Fig. 3F). To confirm this, we estimated half-lives of each gene from FISH spot densities measured after 0, 6, and 22 hr of transcriptional inhibition by actinomycin D. We used these half-lives to predict the fraction of cytoplasmic transcripts that should remain after 18 hr of nocodazole treatment if new transcripts were unable to exit the nuclear periphery (Fig. S5A and S5B). The predictions of this decay model correlated well with observations (Pearson R = 0.71), supporting the idea that RNAs observed in the cytoplasm following culture with nocodazole were already present prior to treatment (Fig. 3G).

Due to its short half-life, nearly all cytoplasmic Polr2a mRNAs decayed during 18 hr of nocodazole treatment; thus, any molecules observed in the cytoplasm after nocodazole washout were highly likely to have traveled there from the perinuclear region after microtubules re-polymerized. We allowed a sparse microtubule network to re-polymerize after 18 hr nocodazole treatment by a partial washout that left trace amounts of nocodazole in the culture medium. We found Polr2a mRNAs in the cytoplasm only at the ends of microtubule filaments connected to nuclei (Fig. 3H), indicating that these are the only routes of escape for mRNAs to leave the perinuclear region. After 30 min, few RNAs were present at the ends of these segments, but substantial perinuclear accumulation remained, suggesting that egress of RNPs along microtubules is a rate-limiting step.

Interestingly, we also found that RNA density for all genes except Ttn and Myom1 increased during 18 hr of culture without treatment as compared to fibers fixed immediately after isolation (Fig S5C). Expression of Hist1h1c, which exhibited the strongest upregulation, was also noticeably heterogeneous between nuclei (Fig. S4A and Video S3). Similar heterogeneous bursts of transcription following myofiber culture have been observed for Myod1, resulting in localization of mRNA near myonuclei (Kann and Krauss, 2019). Compared to freshly isolated fibers, upregulated RNAs in cultured fibers were noticeably enriched at the myofiber surface and closer to nuclei on average. In the cytoplasm surrounding myonuclei in which mRNAs were highly upregulated, mRNAs localized furthest from progenitor nuclei along the longitudinal nuclear poles, from which the most robust microtubule bundles extend (Fig. S4A and Video S3). Given these observations, we infer that RNPs depart myonuclei most efficiently along microtubule bundles at the myofiber surface, but require additional time to disperse throughout the fiber if recent bursts of transcription have occurred.

### RNA Diffusion Is Increasingly Restricted During Muscle Differentiation

In adult myofibers, RNAs may depend on microtubules to leave the perinuclear region due to the constraints on diffusion imposed by mature sarcomeres (Papadopoulos et al., 2000). To study RNA mobility across different stages of sarcomere maturity, we used the C2C12 myoblast cell line. These myoblasts are mononucleated and mitotically active, but can undergo cell cycle arrest and fusion upon serum withdrawal to form multinucleated myotubes. While cultured C2C12 myotubes do not mature to the level of *ex vivo* myofibers, they can generate sarcomeric structures when cultured on patterned hydrogels (Denes et al., 2019). We examined Polr2a mRNA localization in nocodazole-treated C2C12 myoblasts and myotubes (Fig. 3I). In myoblasts, microtubule depolymerization did not result in perinuclear RNA accumulation, and RNAs were frequently observed at the cell periphery. In myotubes, however, microtubule depolymerization did restrict Polr2a mRNA localization near nuclei, but not to the extent observed in adult myofibers. These observations suggest that diffusion of RNAs becomes increasingly limited as sarcomere formation progresses.

### RNAs Trapped in the Perinuclear Region Gather in Large Granules and Inhibit Nuclear Export

We noticed that some RNAs trapped in the perinuclear region after microtubule depolymerization formed large “blobs” rather than individual diffraction-limited spots (Fig. 4A). We quantified the intensities of FISH blobs for each gene in each condition (Fig. 4B) and focused on the most intense blobs (top 5%). For four of the eight genes examined, blob intensities were significantly increased in the perinuclear region (but not cytoplasm) following nocodazole treatment (p < 0.05 by 95% CI overlap, Fig. 4C and 4D). After removing nocodazole for 4 hr, intensities of these large blobs decreased to levels at or below those observed in untreated myofibers. Dmd was a notable outlier: its largest perinuclear granules decreased in intensity in response to nocodazole treatment and remained diminished after washout.

**Figure 4.**
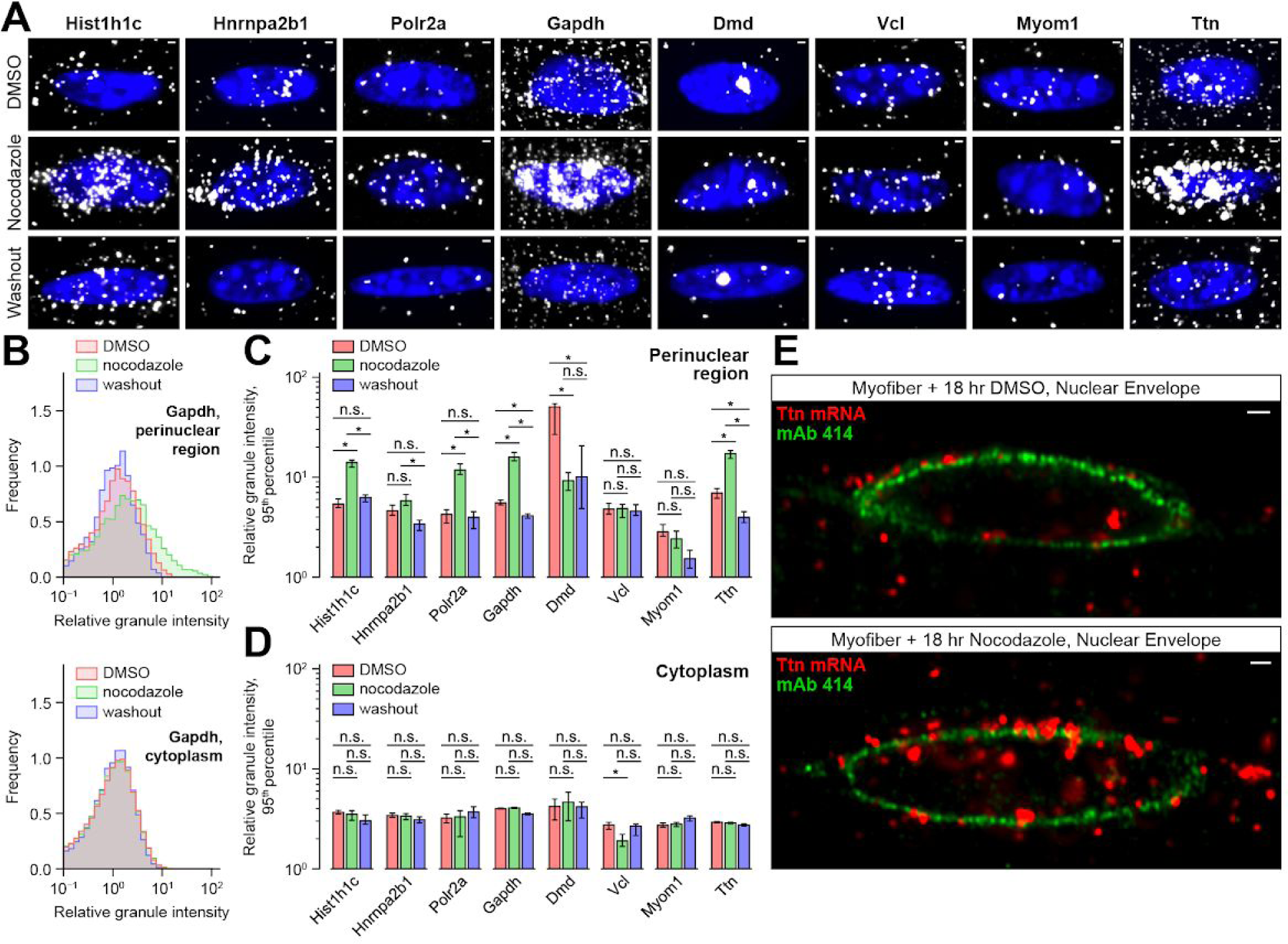
RNAs Trapped in the Perinuclear Region Gather in Large Granules and Inhibit Nuclear Export. A) Representative zoomed-in regions of individual nuclei from the myofiber FISH images described in Fig 3. Scale bars: 1 µm. B) Distribution of Gapdh mRNA blob intensities from myofiber FISH images described in Fig 3. Intensities were normalized to the median cytoplasmic blob within each image. C) The 95th percentile intensity for perinuclear FISH blobs in control, nocodazole-treated, and nocodazole washed-out fibers for each gene. Blob intensities were quantified as in (B) for each gene studied. Error bars = 95% CI. n.s.: not significant, *p<0.05, by confidence interval overlap. Confidence intervals were estimated by bootstrapping. D) The 95th percentile intensity for cytoplasmic plots in control, nocodazole-treated, and nocodazole washed-out fibers for each RNA. Error bars = 95% CI. n.s.: not significant, *p<0.05, by confidence interval overlap. Confidence intervals were estimated by bootstrapping. E) Axial optical cross-section through center of myonucleus in myofiber treated with nocodazole (right) or DMSO (left) for 18 hr and co-labeled by IF/FISH for Ttn mRNA (red) and nuclear pore complex proteins (mAb 414, green). Scale bars: 1 µm.

We found that large intranuclear FISH blobs at putative transcription sites were reduced after nocodazole treatment and did not recover after washout, suggesting transcriptional feedback in response to transport blockade (Fig. 3C, 3D, 4A, and S4A). This transcriptional downregulation would explain reduced blob intensities after washout as compared to untreated fibers. This may also explain the reduced perinuclear blob size for Dmd after nocodazole treatment: due to technical limitations, it is possible that remarkably intense blobs that were actually Dmd transcription sites were mis-classified as perinuclear, and thus the reduction in perinuclear blob intensity in response to nocodazole may actually reflect downregulation of transcription in this case. This technical issue did not impact other genes, as Dmd was the only gene with exceptionally large blobs in the untreated condition.

We additionally observed accumulation of RNAs at the intranuclear periphery after nocodazole treatment (Fig. 3C, 3D, and S4A). To better resolve mRNAs around the nuclear envelope, we co-labeled nuclear pore complex proteins and Titin mRNA in nocodazole-treated and untreated myofibers. We observed RNAs accumulating on both faces of the nuclear envelope and within the nuclear pore, suggesting inhibition of nuclear export (Fig. 4E). Together, these observations suggest that inhibiting RNP transport along microtubules induces formation of large perinuclear RNA granules, potentially causing nuclear export defects and transcriptional inhibition.

### Spatially Dispersed Translation in the Cytoplasm Requires Intact Microtubules

As all mRNAs we studied encode proteins, we investigated the impact of homeostatic and perturbed RNA distribution on protein synthesis. We used HCR RNA FISH to label the ribosomal large subunit rRNA and *in situ* puromycylation to label sites of protein synthesis (Fig. 5A-5C and S6A-S6D). Consistent with previous studies in cardiomyocytes, rRNA was localized along Z-disks in unperturbed conditions (Rudolph et al., 2019). We also observed prominent rRNA signal along microtubule bundles and in the perinuclear region (Fig. 5A and S6A). Ribosomes outside the perinuclear region were evenly dispersed along the length of the myofiber, similarly to mRNA; in contrast to mRNAs, ribosomes were highly concentrated at the myofiber surface (Fig. S6B). Puromycylation signal was nearly identical to rRNA FISH: nascent protein synthesis was localized both within the perinuclear region and along Z-disks and microtubules throughout the length of the fiber, but concentrated near the surface (Fig. 5B, 5C, S6C, and S6D).

**Figure 5.**
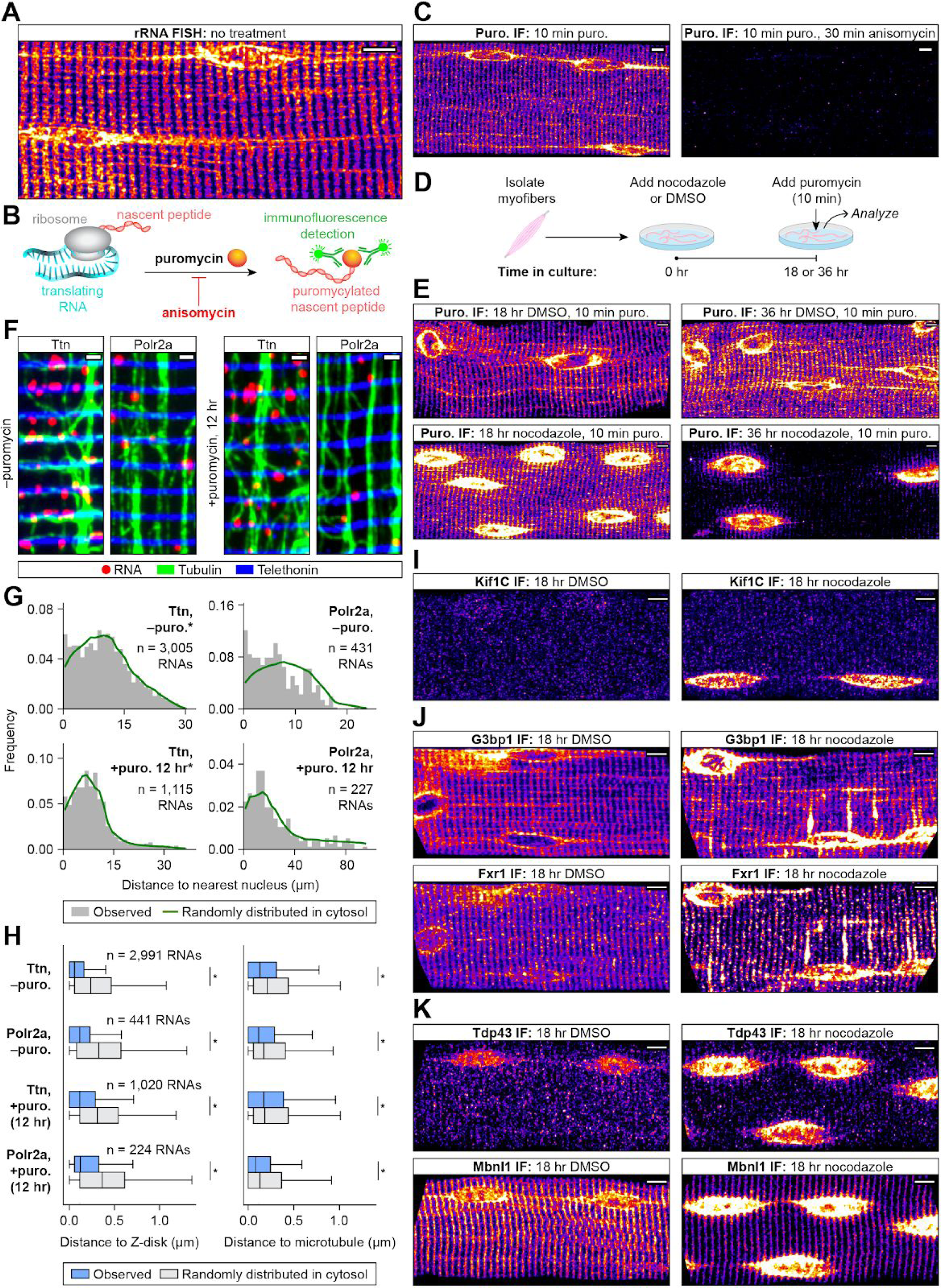
Spatially Dispersed Translation in the Cytoplasm Requires Microtubules, but mRNAs Are Localized Independently of Translation. A) FISH of 28S ribosomal RNA in an isolated myofiber. rRNA signal localized in the perinuclear region and along Z-disks and microtubule bundles, but was concentrated near the myofiber surface. Scale bars: 5 µm. See also Fig. S6A and S6B. B) Schematic describing *in situ* puromycylation to detect nascent proteins in isolated myofibers. Puromycin (2 μM) was added to cultured myofibers for 10 min, producing puromycylated nascent peptides detected using anti-puromycin monoclonal antibodies. Control myofibers were pre-treated with 100 μM anisomycin for 30 min to prevent puromycin incorporation. C) IF against puromycin in a myofiber labeled with puromycin or control as described in (B). Puromycylation signal was nearly identical to rRNA: in the perinuclear region and along Z-disks and microtubules, but concentrated near the myofiber surface. Scale bars: 5 µm. See also Fig. S6C and S6D. D) Live myofibers were isolated and cultured with nocodazole or DMSO (control) for 18 or 36 hr. At each time point, *in situ* puromycylation was performed as described in (B). E) IF against puromycin in myofibers from the experimental time course described in (D). Puromycylation signal was elevated in the perinuclear region, absent from microtubules, and remained along Z-disks in myofibers treated with nocodazole for 18 hr. Puromycylation was confined to the perinuclear region and nearly absent from the cytoplasm in myofibers treated with nocodazole for 36 hr. Scale bars: 5 µm. See also Fig. S6E. F) IF/FISH co-labeling of Ttn or Polr2a mRNAs (red), telethonin protein (Z-disks, blue), and tubulin protein (microtubules, green) in untreated myofiber (from Fig. 2A, left) and in a myofiber treated with 100 µM puromycin for 12 hr to inhibit translation (right). Scale bars: 1 µm. See also Fig. S6F. G) Distance to nucleus for Ttn and Polr2a mRNAs (grey bars) measured in untreated myofibers (from Fig. 1F, left) and in myofibers treated for 12 hr with 100 µM puromycin (right). Distance distributions were compared to null distributions generated from random cytoplasmic coordinates (green). Distributions derived from images of at least two FISH-labeled myofibers per gene/condition combination. Shift in median distance to nucleus was similar in puromycin-treated and untreated myofibers. *p<0.05 with shift in median distance >0.5 µm, Mann-Whitney *U* test. H) Distribution of distances from cytoplasmic Ttn and Polr2a mRNAs (blue boxes) to Z-disks (left) and microtubules (right) in untreated (from Fig. 2D, top) and puromycin-treated myofibers (bottom), compared to null distributions generated from randomly selected cytoplasmic coordinates (grey boxes). Data were derived from images of at least two FISH labeled myofibers per gene/condition combination. *p < 0.03, Mann-Whitney *U* test. I) IF against Kif1C protein in a nocodazole-treated or control myofiber. Scale bars: 5 µm. J) IF against G3bp1 and Fxr1 proteins in a nocodazole-treated or control myofiber. Scale bars: 5 µm. See also Fig. S6G, S6H. K) IF against Tdp43 and Mbnl1 proteins in a nocodazole-treated or control myofiber. Scale bars: 5 µm.

To determine whether spatial patterns of ribosomes and protein synthesis are also dependent on microtubules, we performed *in situ* puromycylation and rRNA FISH after 18 hr and 36 hr nocodazole and control treatments (Fig. 5E and S6E). After 18 hr, both rRNA and puromycylation signals were elevated in the perinuclear region and lost from the microtubules relative to control, though enrichment at Z-disks was unchanged. At 36 hr, both signals were highly enriched in the perinuclear region and nearly absent from the cytoplasm. These results suggest that spatially dispersed translation requires microtubules, and that mRNAs trapped in the perinuclear region continue to be translated. The initial persistence of translation in the cytoplasm following 18 hr of nocodazole treatment aligns with our earlier observation that many long-lived RNAs are still present in the cytoplasm in significant numbers at that time point.

Recent studies showed that puromycylated peptides are released and diffuse quickly from the site of translation in cultured cell lines, leading to inaccuracy in identification of translation sites (Enam et al., 2020; Hobson et al., 2020). It is likely that the dense environment of the myofiber restricts diffusion of labeled peptides away from the site of synthesis (Papadopoulos et al., 2000), avoiding the pitfalls of this technique observed in dividing cells.

### mRNAs Are Localized To Z-disks and Microtubules Independently of Translation

It is possible that actively translating ribosomes could recruit mRNAs to the cytoskeleton, or mRNAs could be associated with the cytoskeleton independently of translation. To distinguish between these possibilities, we cultured myofibers for 12 hr in the presence of puromycin—here for long term translation inhibition, rather than label nascent peptides—and we examined the resulting localization patterns for Polr2a and Ttn RNAs (Fig. 5F and S6F). RNAs from both genes remained dispersed throughout the myofiber and associated with Z-disks and microtubules (Fig. 5G and 5H). These results suggest that while the localization of ribosomes and protein synthesis requires microtubules, mRNA localization and association with the cytoskeleton is not dependent on translation.

### Factors Regulating Translation and Transport of RNPs Co-Accumulate With Trapped mRNA

Given that RNAs and nascent protein synthesis require microtubules for efficient localization in muscle, we asked whether the spatial patterns of protein regulators of RNP formation, transport, and translation are also dependent on intact microtubules. We cultured myofibers for 18 hr with nocodazole and stained for a variety of factors by IF. Kif1C is a kinesin motor protein that has been shown to transport mRNAs on microtubules and is highly expressed in muscle (Pichon et al., 2020). We found that Kif1C was diffusely localized in untreated myofibers and accumulated in the perinuclear region following culture with nocodazole (Fig. 5I). Fxr1 and G3bp1 are RBPs that regulate translation (Sahoo et al., 2018; Vasudevan and Steitz, 2007), and mutations in Fxr1 are linked to congenital multi-minicore myopathy (Estañ et al., 2019). Both proteins were found to localize along Z-disks and microtubules, and following nocodazole treatment, both accumulated in the perinuclear region and on Z-disks adjacent to some nuclei (Fig. 5J). We found mRNAs accumulating on nucleus-adjacent Z-disks following nocodazole treatment as well. Upon closer inspection, we found that Z-disks exhibiting accumulation of Fxr1 and G3bp1 or of mRNAs were actually connected to nuclei by nocodazole-resistant microtubule segments that facilitated perinuclear egress (Fig. S6G and S6H). Given these observations, we conclude that RNPs may be able to move within Z-disks but not between them or away from the perinuclear region without microtubules, as we never observed accumulation of these molecules on Z-disks that did not have a microtubule connection to a nucleus.

Tdp-43 and Mbnl1 are RBPs that shuttle between the nucleus and cytoplasm. Both are known to regulate RNA splicing (Arnold et al., 2013; Kanadia et al., 2003) and localization (Chu et al., 2019; Wang et al., 2012), and are associated with neurological and muscle disease (Thornton, 2014; Weihl et al., 2008). In untreated myofibers, we found that Tdp-43 was enriched in the nucleus, and Mbnl1 showed strong association with Z-disks. These distinct localization patterns suggest differences in the way these RBPs may regulate RNA localization in myofibers. Both RBPs became highly concentrated in the perinuclear region following culture with nocodazole (Fig. 5K). Together, these results illustrate that many protein factors involved in mRNP granule formation, transport, and translation co-accumulate with mRNAs in the perinuclear region upon microtubule disruption.

### RNAs Exhibit Restricted Diffusion and Directed Transport in Live Myotubes

Our observations suggest that mRNAs in myofibers cannot escape progenitor nuclei by passive diffusion and completely depend on microtubules to move throughout the cell. However, we inferred these conclusions via pharmacological inhibition of transport machinery followed by fixation and imaging, and not by observing RNAs in real-time. To confirm inferences made by analyses in fixed myofibers, we used the MS2 system to fluorescently label mRNAs in live myotubes differentiated from C2C12 myoblasts (Fig. 6A) (Buxbaum et al., 2015). We generated a stable C2C12 myoblast line expressing MS2 coat protein (MCP) fused to HaloTag for visualization, a ponasterone-inducible Kdm5b reporter RNA with 45x MS2 hairpins in the 3’ UTR, and a ponasterone-sensitive transcriptional activator. With this cell line, we were able to successfully image single RNA molecules in live myotubes.

**Figure 6.**
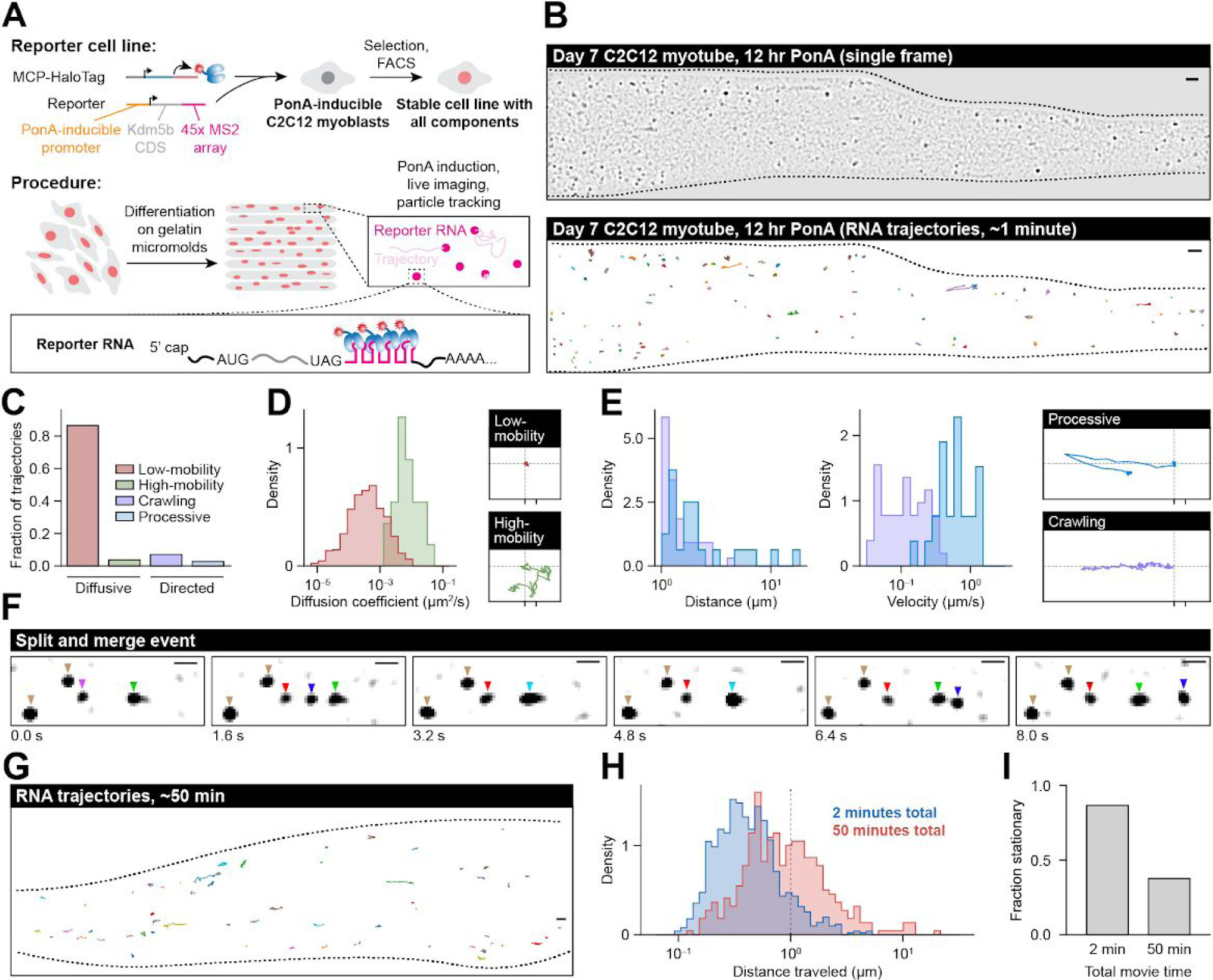
RNAs Exhibit Restricted Diffusion and Directed Transport in Myotubes. A) Schematic describing MS2 live cell RNA imaging strategy. B) Example image of a myotube expressing MS2-labeled RNA (top) and RNA trajectories (bottom) from a 53-second imaging time course. Scale bars: 2 µm. See also Video S4. C) Trajectories were categorized according to their motion (see Methods). See also Video S5. D) Diffusion coefficients calculated for “low mobility” (red) and “high mobility” (green) diffusive tracks (left, see Methods), and example tracks from both groups (right). X-axis ticks for scale: 0.5 µm. E) Distance (left) and velocity (right) calculated for “processive” runs (blue) and “crawling” trajectories (purple, see Methods), and example tracks from both groups (far right). X-axis ticks for scale: 0.5 µm. F) Series of images showing RNA particle splitting and merging events from movies analyzed in A-E. Colored triangles denote particle identities. Scale bars: 1 µm. See also Video S6. G) Example RNA tracks from a myotube expressing MS2-labeled RNA during a 50-minute imaging time course. Scale bars: 2 µm. See also Video S7. H) Maximum distance traveled for each track from the 50-minute (red) and 2-minute (blue) imaging time courses. Tracks below dotted line (1 µm) are categorized as stationary. I) Fraction of stationary tracks in 50-minute and 2-minute time courses.

We imaged the MS2 reporter RNA continuously, at frame rates >1 per second, to characterize particle motion dynamics. We observed individual puncta throughout the cytoplasm, and we tracked the motion of particles in five movies to obtain 699 individual mRNA trajectories (Fig. 6B and Video S4). After careful observation, we categorized particles into four motion states based on their motion—two diffusive and two directed (Fig. 6C and Video S5). Within the diffusion category, we classified particles that moved <1 μm throughout the entire time course as “low-mobility” (605 trajectories, 86%). Most RNAs in the “low-mobility” state appeared truly stationary (median diffusion coefficient 3.9 x 10^-4^ µm^2^/s), and at the high end of the distribution, their diffusion coefficients resembled what has been observed in neurons (max. diffusion coefficient 9.0 x 10^-3^ µm^2^/s, versus 3.8 x 10^-3^ µm^2^/s in neurons) (Park et al., 2014), which is an order of magnitude lower than RNA diffusivity in fibroblasts (Fig. 6D) (Park et al., 2014). A small subset of RNAs moved >1 μm in a non-directed fashion (26 trajectories, 4%), and we classified these RNAs as “high-mobility” (Fig. 6D). The most mobile of these particles were nearly as diffusive as RNAs in fibroblasts (max. diffusion coefficient 0.04 µm^2^/s, versus 0.09 µm^2^/s in fibroblasts) (Park et al., 2014), but they were rare and typically localized in clusters of “high-mobility” RNAs at the myotube periphery. After closer inspection, we believe that this state is an artifact of studying RNA dynamics in immature C2C12 myotubes, and that “high-mobility” RNAs occur only in underdeveloped regions of myotubes that lack sarcomere structures, which are not representative of mature myofibers.

“Directed” particles were defined as those that moved >1 μm in a single direction (68 trajectories, 10%). We further classified these RNAs as “processive” if they took at least one contiguous run >1 μm, or “crawling” otherwise (Fig. 6E). “Processive” runs exhibited behavior characteristic of directed transport of RNP granules along microtubules by motor proteins such as kinesin and dynein, and have been observed in a variety of other cell types (Liao et al., 2019; Park et al., 2014; Turner-Bridger et al., 2018). Overall run velocity was slightly slower (median 0.63 μm/s) than velocities typically observed in neurons (∼1 μm/s), although the fastest runs did reach almost 1.5 μm/s. Most runs were short (median run length 1.6 μm), but the distribution of travel distances had a long tail with multiple runs of tens of microns observed (max. run length 17 μm). The slow, prolonged “crawling” motion that we observed is to our knowledge a novel transport state for RNAs. Speeds in this state were slower than “processive” runs (median 0.11 μm/s), and although the median distance traveled by these particles was similar to median “processive” run length (1.11 μm), the tail of the distance distribution was much shorter (max. distance 4 μm). Together, these data reinforce the conclusions drawn in *ex vivo* myofibers: RNA diffusivity is highly restricted in muscle, and directed transport is prominent. RNP transport granules have been shown to merge and split in other cell types, and we observed multiple examples of these behaviors in myotubes as well, indicating that these mRNPs can form higher order granules or complexes (Fig. 6F and Video S6).

By imaging with relatively fast frame rates, these particle tracking experiments provided detailed information about the motion of RNAs over ∼1 min, and showed that most RNAs remain nearly stationary on this time scale. To determine whether RNAs eventually move from these locations, we performed a longer imaging time course in which we captured frames every 15 s for 50 min (Video S7). We tracked 367 individual particles in four movies (Fig. 6G), and found that 141 out of 367 RNAs (38%) still moved <1 μm over the entire 50 minutes (Fig. 6H and 6I), indicating that the fraction of particles undergoing motion did not scale linearly with time. This runs contrary to observations in neurons, where directed events reposition RNAs at a constant rate (Yoon et al., 2016). This indicates that some cytoplasmic particles in myotubes may be permanently stationary once anchored, while others are more competent for directed motion.

### Computational Simulation Confirms that Directed Transport is Required for Near-uniform Localization of mRNA in Myofibers

After obtaining quantitative motion parameters from live-cell imaging of C2C12 myotubes, we integrated these measurements into a 3D stochastic simulation of RNA transport in myofibers. In our discrete-time Markov chain (DTMC) model, RNAs are generated at the periphery of myonuclei and travel throughout the cytoplasm until they decay (Fig. 7A). While traversing the fiber, RNAs are assigned to a set of possible motion states, including diffusion and directed transport. RNAs transition between these states randomly such that the average number of granules in each state matches the expected distribution from C2C12 myotube imaging (Fig. 7B). Diffusion coefficients and distances traveled during directed transport events are sampled from distributions estimated from live-cell imaging measurements, and production and decay rates were constrained by observed RNA copy numbers and half-lives (Fig. 1D and S5B). Using fiber and nuclei segmentations obtained from confocal images, we applied this model to simulate RNA transport dynamics within the geometry of real myofibers.

**Figure 7.**
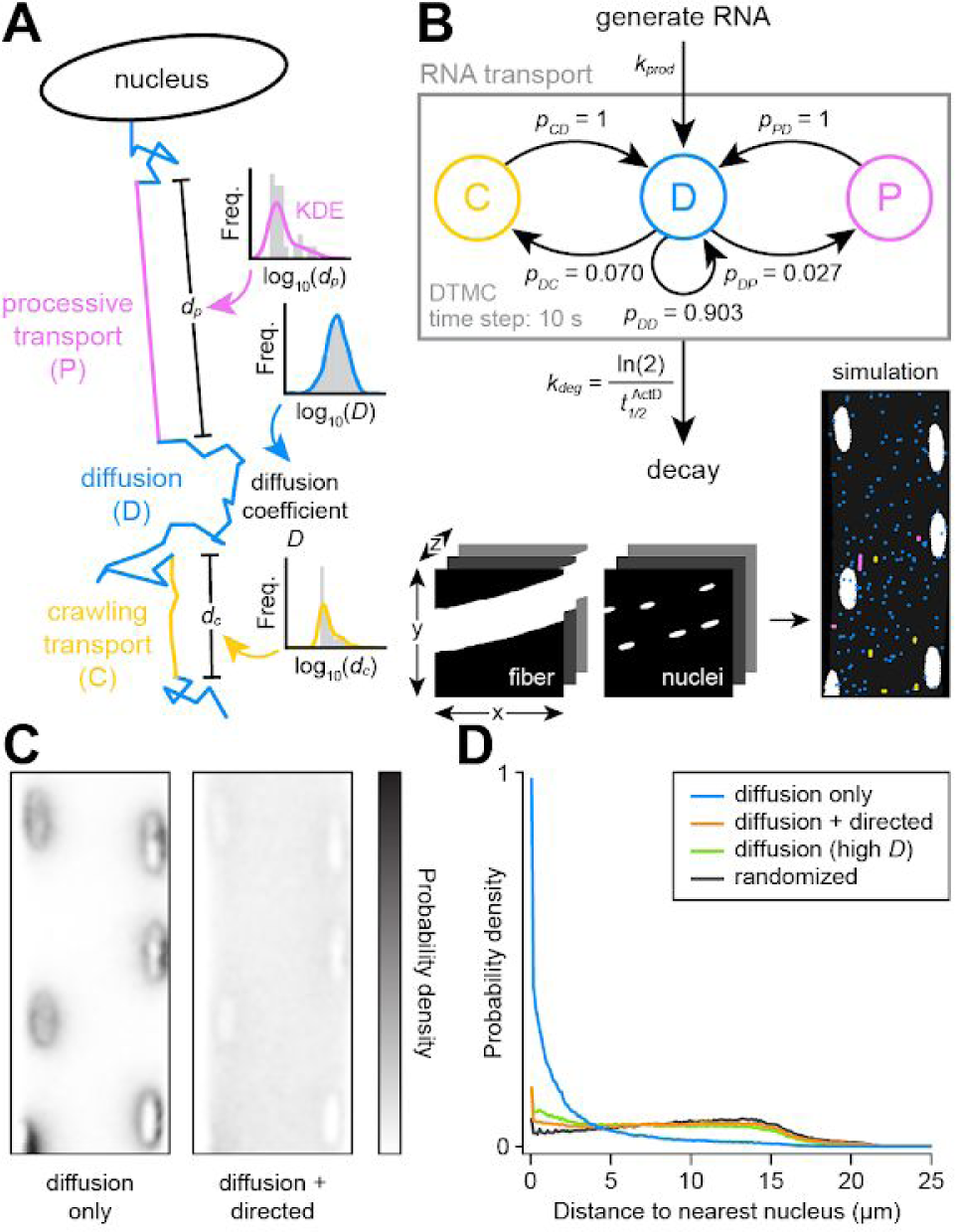
Computational Simulation Confirms That Directed Transport Is Required to Disperse mRNA in Myofibers. A) Diagram of RNA mobility states modeled in the simulation of RNA motion in myofibers. RNA molecules were spawned from the nuclear envelope and allowed to transition between “low mobility” diffusion and active transport states. While in each state, RNAs moved in 3D space according to parameters sampled from distributions shown in Fig. 6D and 6E obtained from C2C12 myotube imaging. B) Network diagram of the discrete-time Markov chain (DTMC) model developed to simulate RNA generation, motion, and decay. Production and decay rates were constrained by measured mRNA density and half-life (See Fig. 1 and Fig. S5). Transition probabilities were derived from the average number of RNAs observed in each state in C2C12 myotubes (See Fig. 6C). Simulations were conducted in 3D using myofiber and nuclear geometries segmented from confocal images. C) Comparison of RNA localization patterns observed in 1000 hr simulations of Polr2a RNA either with (right) or without (left) directed transport states. The average RNA density was calculated by sampling once every hour, with the first 10% of the simulation discarded as burn-in. See also Video S8. D) Distance to nucleus measured for simulated Polr2a mRNAs (see C) either with (orange) or without (blue) directed transport states. Shown for comparison is the distribution from a simulation of Polr2a mRNAs in the “high mobility” diffusion state (green, see Fig. 6D) and a null distribution generated from randomly selected cytoplasmic coordinates (grey).

We first simulated transport of Polr2a mRNA under the constraint that RNAs only move in the “low-mobility” passive diffusion state. In this configuration, in the absence of directed transport, we found that the steady-state distribution showed strong accumulation of RNA around nuclei and depletion in nucleus-distal regions (Fig. 7C, 7D, and Video S8), with 58% of RNAs observed <2 μm from a nucleus. This profound perinuclear accumulation starkly contrasts our experimental observations of Polr2a RNA in untreated myofibers. However, when we allowed RNAs in the simulation to transition between diffusion and active transport states, including both “processive” and “crawling” states, we found that the distribution of RNA in the cytoplasm approached uniformity at equilibrium. When compared to a uniform randomized null distribution, we still observed a slight perinuclear accumulation of RNAs (14% within 2 μm from a nucleus, compared to 8% for randomly distributed RNA), similar to what we observed experimentally for several genes in fixed myofibers. This likely results from the requirement of an initial directed transport event to exit the perinuclear region.

As a comparison, we also performed simulations of Polr2a mRNA in the “high-mobility” state (Fig. 7D and Video S8), in which the diffusion coefficients are similar to those previously observed in fibroblasts (Park et al., 2014). In this context, we found that the steady-state distribution of RNA indeed approached uniformity and resembled experimental observations (Fig. 7D). Therefore, in principle, if RNPs in myofibers could move as freely as they do in dividing cells in culture, diffusion alone would likely be sufficient for efficient population of nucleus-distal regions. Directed transport observed in muscle thus appears to be an adaptation that facilitates efficient distribution of gene cargoes within a dynamic and densely packed syncytium.

## DISCUSSION

Basic principles of mRNA localization are not well understood in skeletal muscle, and it is unknown whether mechanisms operating in other differentiated cell types also apply to muscle syncytia. Here, we developed a method to detect single RNA molecules and protein markers in mature skeletal muscle fibers, and we characterized spatial patterns of a diverse panel of RNAs in homeostatic and perturbed conditions. All mRNAs that we examined are widely dispersed in the cytoplasm, readily traveling at least tens of microns from progenitor nuclei. We also found that enrichment at sarcomere Z-disks is not limited to sarcomere-encoding mRNAs, but instead is a common feature of all mRNAs we studied. Ribosomes and protein synthesis are also localized at this structure, suggesting a role for the Z-disk as a biosynthesis hub. We discovered that muscle development imposes stark restrictions on passive RNA mobility, and that in adult myofibers, microtubule-dependent directed transport is absolutely essential for RNAs to travel away from progenitor myonuclei. Disrupting mRNA transport leads to accumulation of large RNP granules and nascent protein in the perinuclear region, with potential implications in muscle disease.

The widespread distribution of RNAs may facilitate optimal distribution of protein products, as well as confer robustness to transcriptional bursting. Interestingly, the extent of dispersion from myonuclei was correlated with expression level and not with transcript length, and was not strictly related to encoded protein function (Fig. 1 and S1). Even exceptionally large mRNAs, such as Ttn, travel efficiently in the myofiber. Additionally, mRNAs encoding nuclear proteins are likely often translated in the vicinity of nuclei from which they did not originate, suggesting a potential mechanism for inter-nuclear gene regulation. Recent studies of myofibers revealed enrichment of some RNAs near myonuclei, and proposed that recent bursts of transcription can lead to this localization pattern (Kann and Krauss, 2019). Our observations of mRNAs localizing closer to nuclei following culture-induced upregulation supports this interpretation (Fig. 3, S4 and S5).

We were surprised to find that all mRNAs studied are localized along sarcomere Z-disks (Fig. 2 and S2). Decades ago, Z-disk RNA localization was proposed to play a role in sarcomere assembly (Fulton, 1993). However, mRNAs encoding nuclear, metabolic, and even M-line proteins are enriched at Z-disks, together with ribosomes and global protein synthesis machinery (Fig. 5 and S6). These observations evoke a model in which proteins are translated at the Z-disk, yet exhibit sufficient mobility to travel to their final destinations. Studies in cardiomyocytes suggest this model may be broadly applicable to striated muscle cells. For example, Ttn mRNA is localized to Z-disks in cardiomyocytes, but Ttn protein is mobile and likely incorporates into the sarcomere as full-length protein from a soluble pool (Rudolph et al., 2019). Protein synthesis and turnover machinery are localized to Z-disks in cardiomyocytes as well (Lewis et al., 2018; Rudolph et al., 2019), and we predict that most mRNAs will localize to Z-disks in these cells. Why and how the Z-disk recruits RNAs to serve as a biosynthesis hub is not fully clear, but endoplasmic and sarcoplasmic reticular membranes are concentrated at Z-disks and may play a role (Villa et al., 1993). Similarly to other cell types, it is possible that actin filaments or actinin itself directly tether RNPs. Alternatively, during muscle contraction, myosin thick filaments may evict RNPs from the A-band region and concentrate them at Z-disks. Translation of RNAs along microtubules was also observed, but RNA localization did not depend on translation; further work will be required to determine whether translation occurs on RNPs in motion.

Completely unexpected was the absolute dependency on microtubules for RNAs to move more than a few microns from the nucleus (Fig. 3). Microtubule dependence for mRNA distribution to the periphery of cardiomyocytes has been observed (Perhonen et al., 1998), but we find a much more severe effect in adult skeletal myofibers. RNA distribution increasingly relies on microtubules as muscle matures, likely due to increasing organization and density of sarcomeres (Fig. 3I). Interestingly, we see some evidence for limited mobility within, but not between, Z-disks in the myofiber cytoplasm (Fig. S6H); however, our data suggests that escape from the perinuclear region is completely dependent on engagement with microtubule-based transport machinery, e.g. kinesins or dyneins. Transport inhibition leads to the formation of large RNP granules, with potential feedback on nuclear export and transcription, highlighting the importance of efficiently clearing RNAs from the perinuclear region (Fig. 4). The grid-like structure of the microtubule lattice in myofibers, with large bundles of microtubules extending from the longitudinal myonuclear poles, likely plays a critical role in moving cargoes quickly from the perinuclear region and distributing them along Z-disks. Indeed, newly produced RNAs move furthest from myonuclei along these large bundles (Fig. S4).

Why are RNAs trapped so abruptly in the perinuclear region, and what could underlie export and transcriptional feedback? The linker of nucleoskeleton and cytoskeleton (LINC) complex tethers the nucleoskeleton to the microtubule network (Tapley and Starr, 2013), facilitates interactions between the nucleus and molecular motors to position myonuclei (Wilson and Holzbaur, 2012), and is involved in mechanosensitive transcriptional regulation (Alam et al., 2016) and mRNP export (Li and Noegel, 2015). This complex may play a role in coordinating exported mRNAs for transport and regulating feedback in response to mRNP accumulation. Several myopathies, including Emery-Dreifuss muscular dystrophy (Nagano et al., 1996) and limb-girdle muscular dystrophy 1B (Muchir et al., 2000), are caused by mutations to ubiquitously expressed nuclear envelope proteins, highlighting the importance of nuclear envelope function in muscle. Recently, the muscle-specific sk-CIP protein was identified, which regulates myonuclear positioning via interactions with microtubule organizing center (MTOC) proteins and the LINC complex (Liu et al., 2020). The unique structure and function of cytoskeletal and nucleoskeletal proteins may regulate or restrict passive RNP egress from the perinuclear region of myofibers.

These findings hold important implications for muscle development, maintenance, and disease. Efficient RNA distribution and protein translation may play roles in facilitating muscle hypertrophy prior to satellite cell recruitment (McCarthy and Esser, 2007) and in forming new myonuclear domains during regeneration. The strong dependency on directed transport to preserve homeostatic RNP distribution points to vulnerabilities that may underlie myopathies. Amyloid-like “myo-granules’’ containing TDP-43 form in developing and regenerating muscle, while related TDP-43+ RNP aggregates are observed in muscle diseases such as inclusion body myopathy (Vogler et al., 2018). Fxr1 mutations have been shown to cause large aggregates of mutated Fxr1 protein and polyA RNA in congenital multi-minicore myopathy (Estañ et al., 2019). In dystrophin-null settings, the microtubule network is highly disorganized and no longer lattice-like (Oddoux et al., 2019). Centralized myonuclei that occur in dystrophic or non-dystrophic settings may lead to inefficiencies in RNP distribution as a result of the less prominent microtubule lattice in the myofiber core (Folker and Baylies, 2013). The observation that colchicine treatment for gout can occasionally cause myopathy is a long-standing mystery; altered vesicle transport has been proposed to play a role, but our observations suggest a possible contribution from alterations to global RNA and protein distribution (Fernandez et al., 2002).

Finally, observations here may provide broad insights into RNA localization regardless of cell type. RNAs have often been proposed to be either localized or non-localized, with the implication that many non-localized RNAs passively diffuse in the cytoplasm. However, in muscle, it appears that any transit away from the nucleus requires microtubule-dependent transport. Here, the sushi-belt model for RNA distribution (Doyle and Kiebler, 2011) could be the rule rather than the exception—all mRNAs must undergo directed transport by default, with specific localization patterns accomplished via anchoring events. In neurons, RNAs devoid of specific localization elements still undergo directed transport, and it is the directionality and frequency of directed runs that are modulated by specific *cis*-elements (Bauer et al., 2019). In both of these cell types, diffusion coefficients of mRNAs are orders of magnitude lower than in cultured dividing cells, suggesting that directed transport occurs by default and RNP properties confer specificity. In contrast to neurons, the total fraction of RNPs undergoing directed motion at any given time is much lower in myotubes, in which a significant fraction of RNAs could be permanently immobile (Fig. 6). This may reflect a biological need for RNAs to be stationary at Z-disks for translation, perhaps even for days at a time, given the long half-lives of some transcripts (Fig. S5).

Which components of RNPs facilitate directed transport? Effective RNA distribution does not depend on ribosomes (Fig. 5), and all mRNAs studied are polyadenylated. Therefore, core machinery such as the cap-binding complex or poly-A binding protein may play important roles, or perhaps multiple RBPs can engage with microtubule motors. Further elucidation of a potential RNA localization code may reveal additional specificities for certain RNPs, with the underlying constraint that, notwithstanding isoform differences, the same RNA sequences must encode localization signals that function properly across numerous cell types and tissues.

## AUTHOR CONTRIBUTIONS

LTD designed and performed all myofiber and cell culture experiments. LTD and CPK designed and implemented all quantitative image and computational analyses. ETW supervised and helped design all aspects of the study. LTD, CPK, and ETW wrote the manuscript.

## ACKNOWLEDGMENTS

We thank Maury Swanson, Karyn Esser, Andy Berglund, Gary Bassell, Laura Ranum, members of the Wang Lab, and members of the University of Florida Center for Neurogenetics for helpful comments and suggestions. This work was supported by an NSF Graduate Research Fellowship to CPK; and NIH R01NS114253 and a Chan-Zuckerberg Initiative Ben Barres Career Acceleration Award to ETW.

---

**Figure S1.**
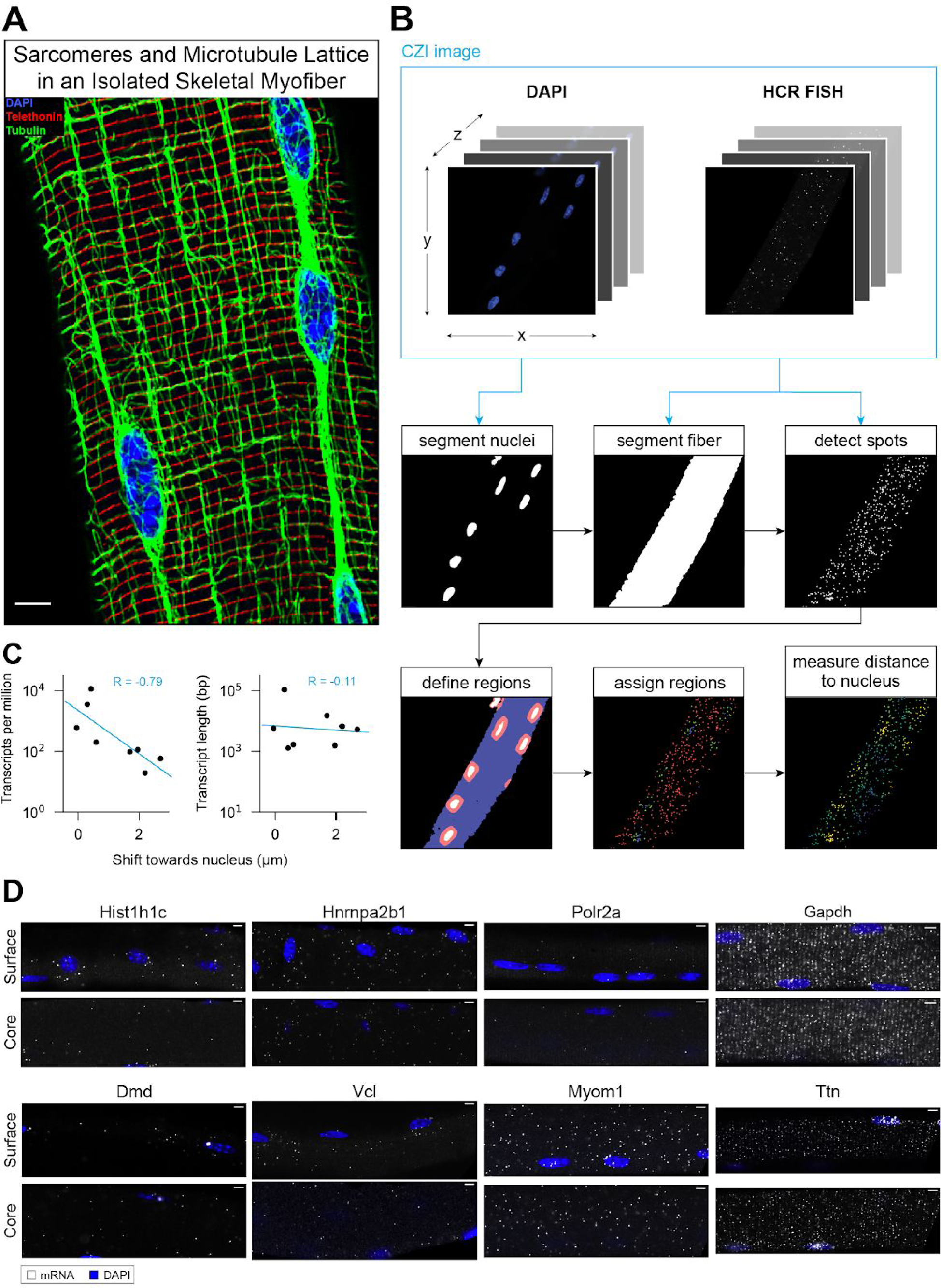
Single mRNAs Can Be Reproducibly and Accurately Localized by HCR smFISH and Are Dispersed Throughout Myofibers, Related to Figure 1. A) Co-IF against tubulin protein (green) to label the microtubule network, and telethonin protein (red) to label the sarcomere Z-disks in an isolated myofiber. Myonuclei labeled with DAPI (blue). Scale bars: 5 µm. B) Schematic describing the computational pipeline used to detect RNA FISH spots and segment myofibers and myonuclei. The distance from each FISH spot to nearest nucleus is measured from coordinates and segmentations. C) The magnitude of the shift towards the nucleus for cytoplasmic mRNAs from each gene studied was inversely correlated to RNA copy number (estimated by RNAseq, left) and uncorrelated with transcript length (right). Trendlines: LLS regression. pearson R=−0.79; p=0.02, Wald test (left). pearson R=−0.11; p=0.79, Wald test (right). D) Single optical sections from the surface and core of myofibers labeled by FISH for each RNA studied. Scale bars: 5 µm.

**Figure S2.**
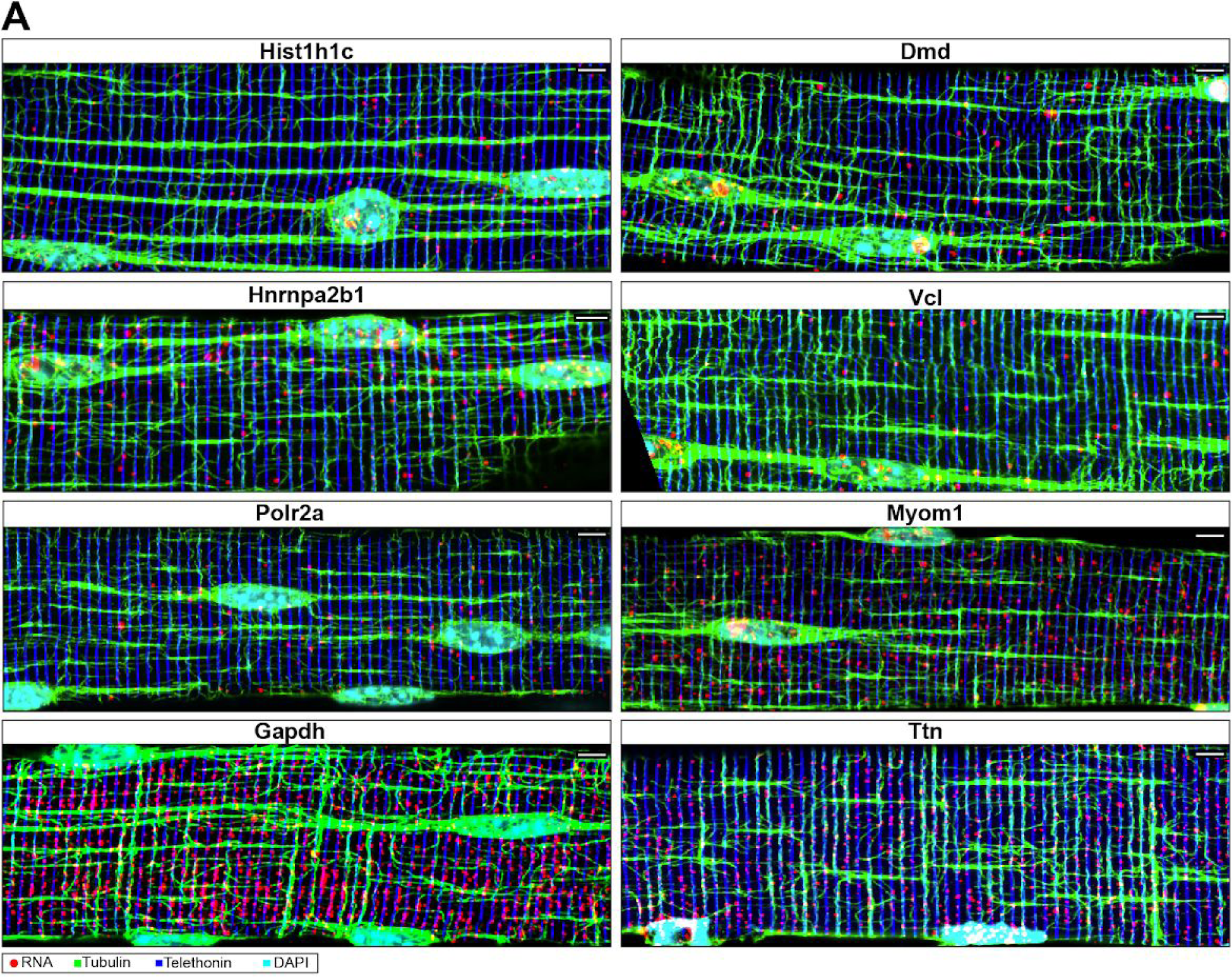
RNAs Associate With Z-disks and Microtubules, Related to Figure 2. A) IF/FISH co-labeling of mRNAs from each gene studied (red), tubulin protein (microtubules, green) and telethonin protein (Z-disks, blue) in isolated myofibers. Scale bar: 5 µm.

**Figure S3.**
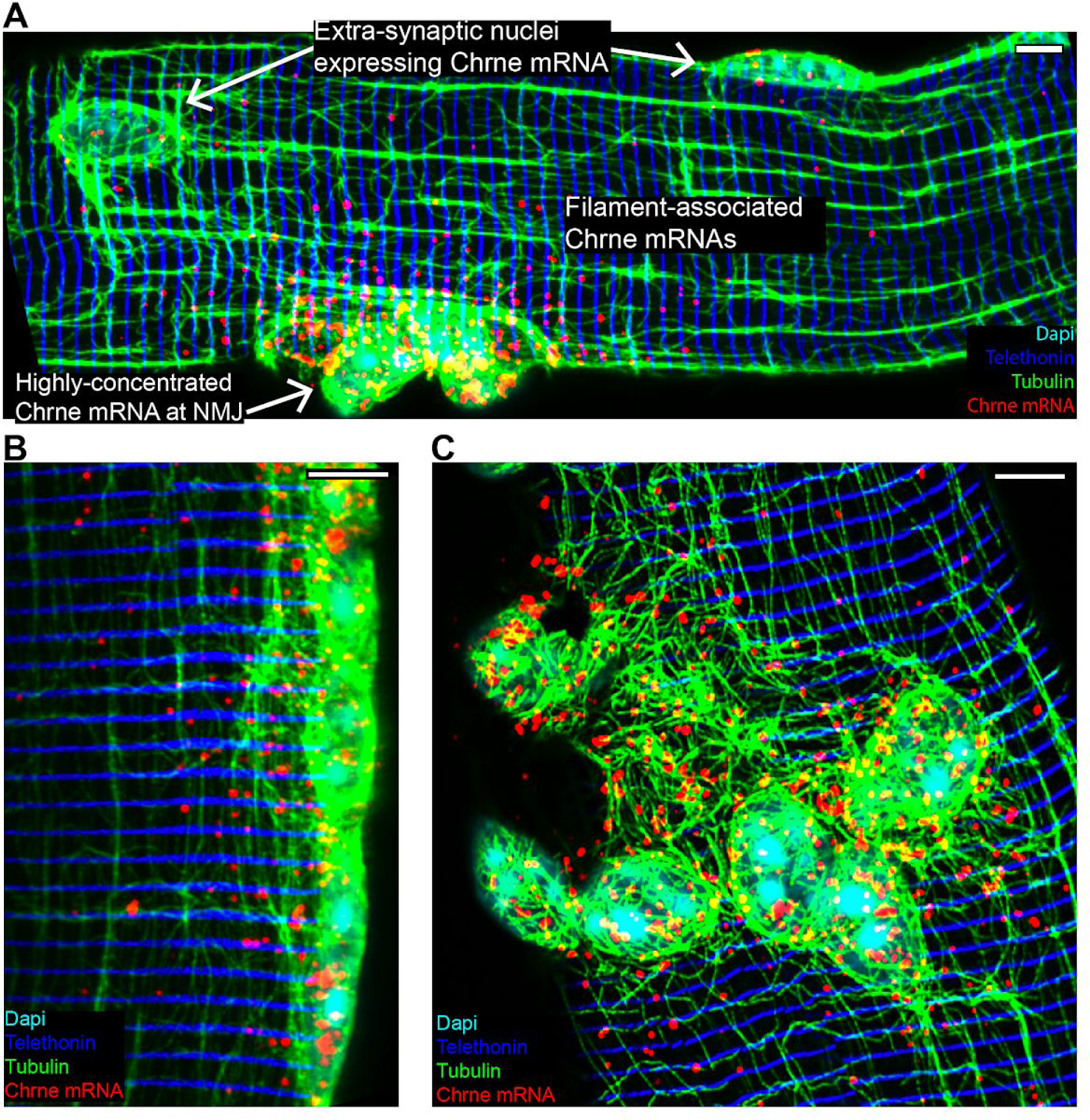
NMJ-Specific RNAs Remain Concentrated at the NMJ, Related to Figure 2. A) IF/FISH co-labeling of Chrne mRNA, tubulin protein (microtubules, green) and telethonin protein (Z-disks, blue) in isolated myofiber. Post-synaptic myonuclei located by characteristic clustered morphology. Scale bars: 5 µm B) Axial optical section of NMJ region in IF/FISH co-labeled myofiber as in A. Scale bars: 5 µm. C) Radial optical section of NMJ in IF/FISH co-labeled myofiber as in A. Scale bars: 5 µm.

**Figure S4.**
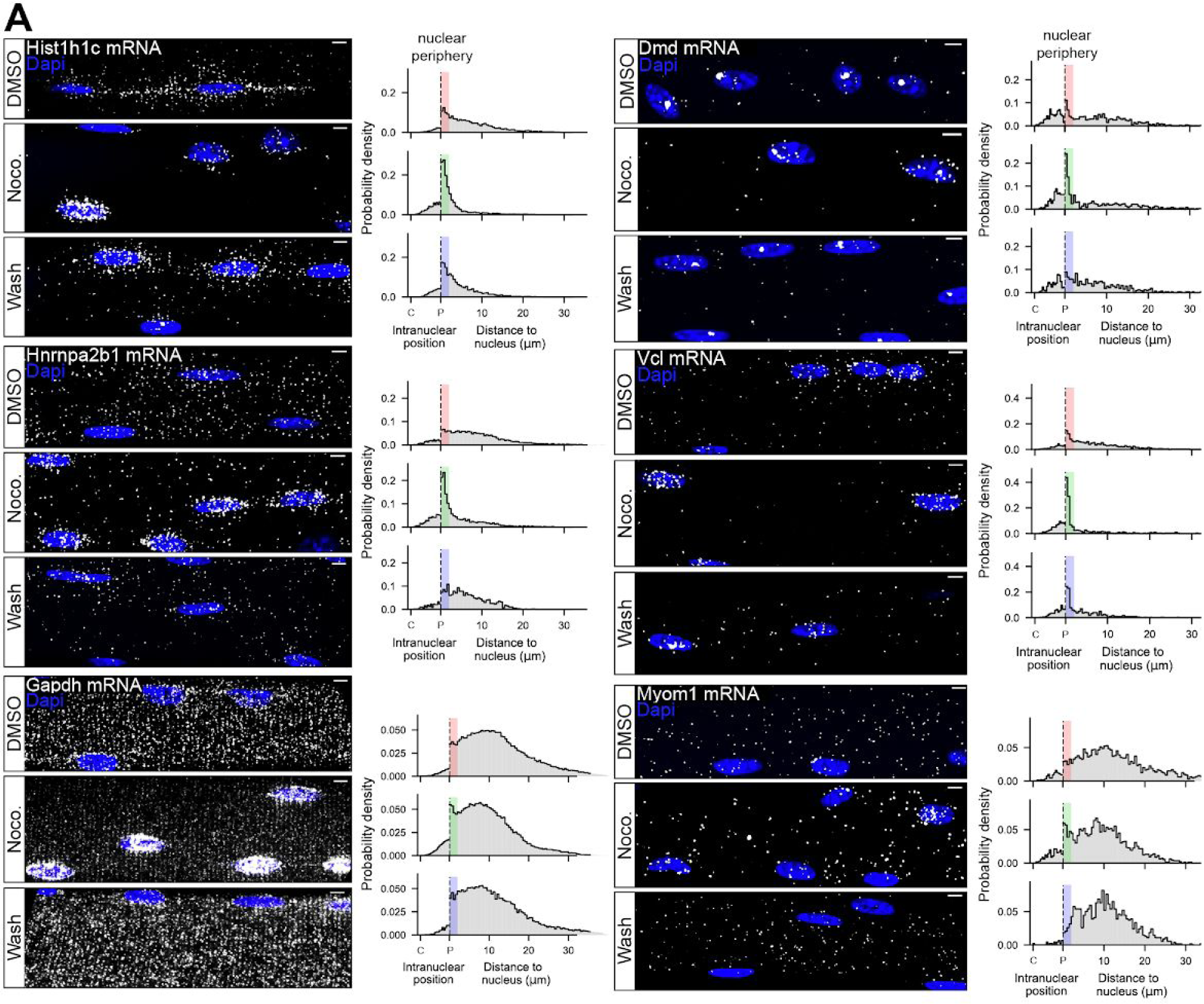
Distribution of RNAs Throughout the Cytoplasm Requires Microtubules, Related to Figure 3. A) FISH of mRNAs from each gene studied in myofibers from the experimental time course described in Fig. 3A (Polr2a and Titin mRNAs shown in Fig. 3C and 3D). Intranuclear position relative to centroid (C) and periphery (P) measured for mRNAs in the nucleus and distance to nearest nucleus measured for mRNAs in the cytoplasm (right). Data derived from multiple images of FISH labeled myofibers in each of two independent experiments. Perinuclear region (defined as 2 μm from the nuclear periphery) analyzed in Fig. 3E denoted by colored bars corresponding to experimental condition. Scale bars: 5 µm.

**Figure S5.**
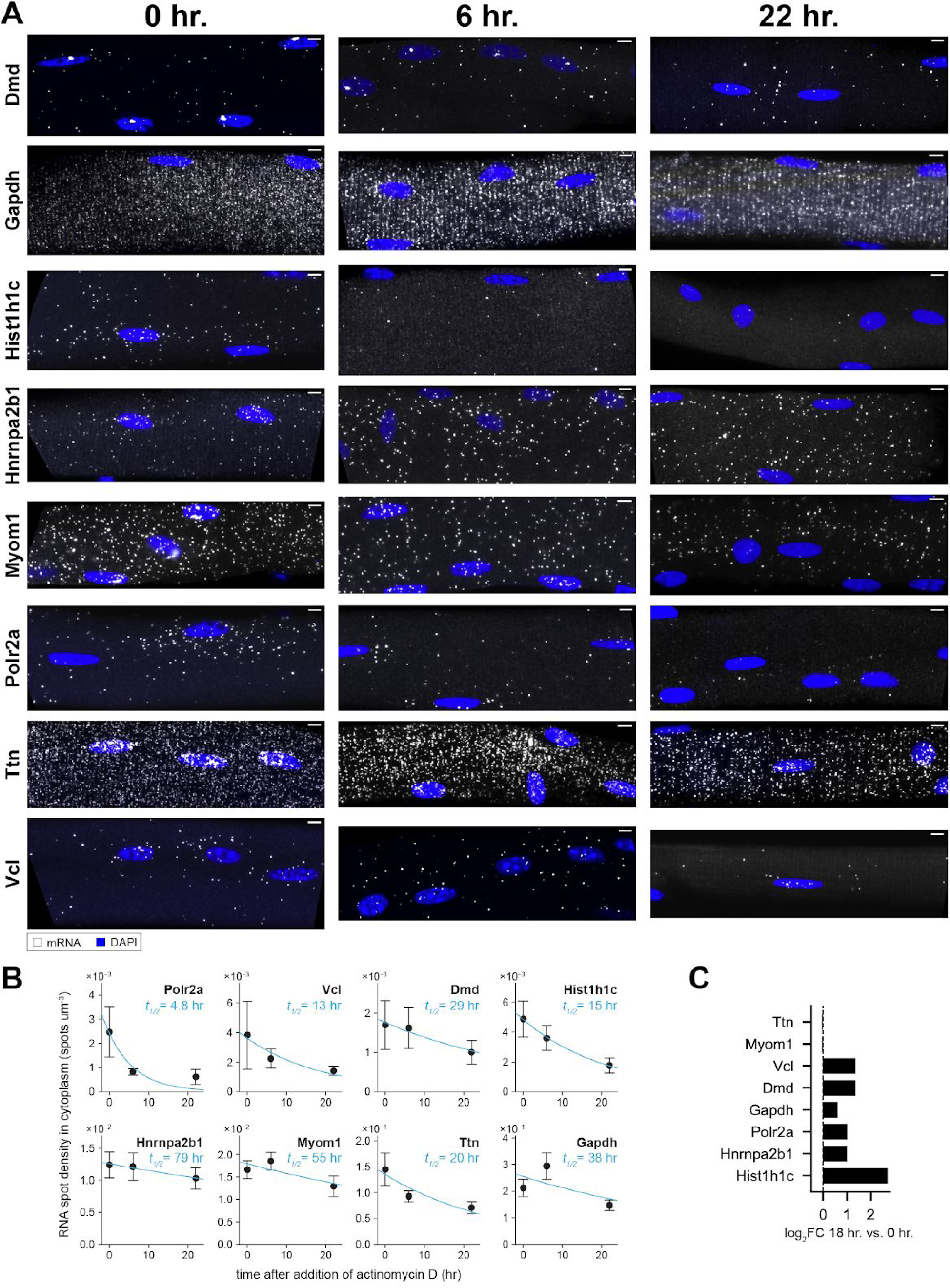
RNA half-lives Predict RNA Distribution After Culture in Nocodazole, Related to Figure 3. A) FISH of mRNAs from each gene studied in myofibers treated with actinomycin D for 0, 6, and 22 hr. Scale bars: 5 µm. B) For each gene studied, mRNA abundances at each actinomycin D time point were measured from images of at least three FISH labeled myofibers. Exponential decay curves were fit to the data to estimate RNA half-lives (t_1/2_). Mean ± SEM. C) Log_2_ fold-change in mRNA density for each gene studied between freshly isolated myofibers and myofibers cultured for 18 hr. Data were derived from at least three images of FISH labeled myofibers per gene/condition combination.

**Figure S6.**
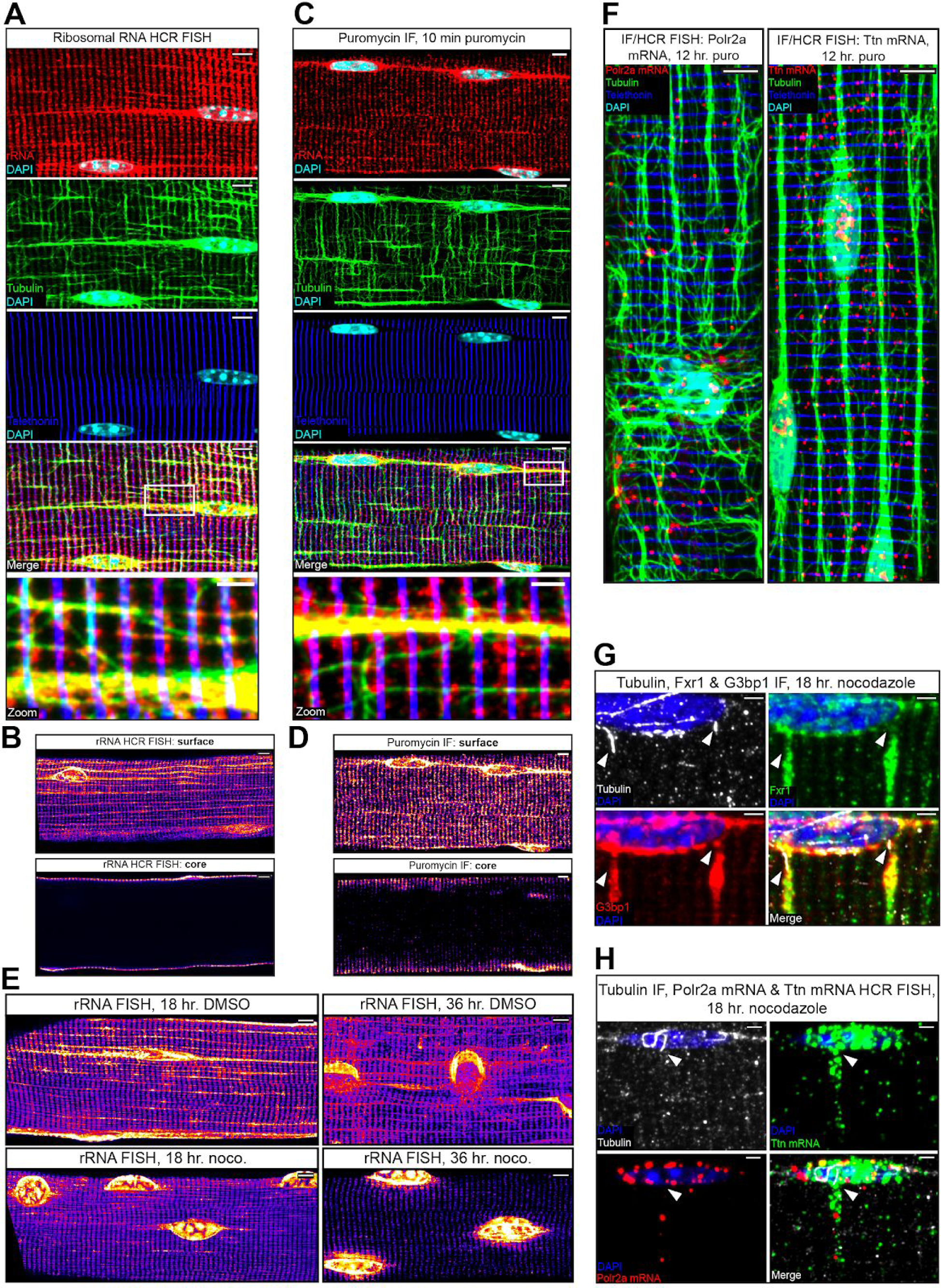
Spatially Dispersed Translation in the Cytoplasm Requires Microtubules, but mRNAs Are Localized Independently of Translation, Related to Figure 5. A) IF/FISH of 28S ribosomal RNA (red), tubulin protein (microtubules, green), and telethonin protein (Z-disks, blue) in a myofiber. Scale bars: 5 µm. B) FISH of 28S ribosomal RNA in axial optical sections of the surface (top) and core (bottom) of a myofiber. Scale bars: 5 µm. C) IF against puromycin (red), tubulin protein (microtubules, green), and telethonin protein (Z-disks, blue) in a myofiber labeled with puromycin or control as described in Fig. 5B. Scale bars: 5 µm. D) IF against puromycin in axial optical sections of the surface (top) and core (bottom) of a myofiber labeled with puromycin or control as described in Fig. 5B. Scale bars: 5 µm. E) FISH of 28S ribosomal RNA during experimental time course analogous to Fig. 5B. rRNA was elevated in the perinuclear region, absent from microtubules, and remained along Z-disks in myofibers treated with nocodazole for 18 hr. rRNA was confined to the perinuclear region and nearly absent from the cytoplasm in myofibers treated with nocodazole for 36 hr. Scale bars: 5 µm. F) IF/FISH co-labeling of Polr2a (left) or Titin (right) mRNAs (red), telethonin protein (Z-disks, blue), and tubulin protein (microtubules, green) in myofibers treated with 100 µm puromycin for 12 hr to inhibit translation (right). Scale bars: 5 µm. G) IF against G3bp1 (red), Fxr1 (green), and tubulin (microtubules, white) in myofibers treated with nocodazole for 18 hr. Accumulation of both RBPs was observed along Z-disks that contacted nucleus-attached, nocodazole-resistant microtubules (arrowheads). Scale bars: 1 µm. H) IF/FISH of Polr2a mRNA (red), Ttn mRNA (green), and tubulin protein (microtubules, white) in myofibers treated with nocodazole for 18 hr. Accumulation of both mRNAs was observed along Z-disks that contacted nucleus-attached, nocodazole-resistant microtubules (arrowhead). Scale bars: 1 µm.

## SUPPLEMENTAL VIDEOS AND LEGENDS

Supplemental videos can be found here: https://data.rc.ufl.edu/pub/ericwang/Denes_et_al_2021/suppvids/

**Video S1. The Microtubule Lattice Interweaves Throughout Sarcomeres of Skeletal Myofibers, Related to Figure 1.** Co-IF of tubulin protein (green) to label the microtubule network and telethonin protein (red) to label the sarcomere Z-disks in an isolated myofiber. Myonuclei labeled with DAPI (blue). Video shows 0.5 µm optical sections extending in 4 µm from the surface of a myofiber. Scale bars: 5 µm.

**Video S2. mRNAs are Localized in the Myofiber Core, Related to Figure 1.** FISH of Ttn mRNA (green) and Hnrnpa2b1 mRNA (red) in isolated myofiber. Myonuclei labeled with DAPI (blue). Video shows 0.5 µm optical sections extending in 8 µm from the surface of a myofiber. Scale bar: 5 µm.

**Video S3. mRNAs Upregulated During Myofiber Culture Localize Furthest from Progenitor Myonuclei Along the Longitudinal Poles, Related to Figure 3.** FISH of Hist1h1c mRNA in an isolated myofiber cultured for 18 hr with DMSO (control treatment, see Fig. S4). Myonuclei labeled with DAPI (blue). Video shows 0.5 µm optical sections extending in 7 µm from the surface of a myofiber. Scale bar: 5 µm.

**Video S4. MS2 Reporter RNAs in Live C2C12 Myotubes, Related to Figure 6.** Videos show five myotubes expressing MS2 labeled RNA (top) with RNA trajectories overlaid (bottom). Scale bars: 2 µm.

**Video S5. Example Tracks from Each Category of RNA Motion. Related to Figure 6.** Particles from myotubes from Video S3 were categorized into one of four motion states (see Methods), two diffusive (“Low Mobility” and “High Mobility”) and two directed (“Crawling” and “Processive”). Scale bars: 0.5 µm.

**Video S6. RNPs Split and Merge in Myotubes, Related to Figure 6.** Zoomed regions of myotubes from Video S2 showing RNPs undergoing directed transport events in which they split from (first video), or split and merge with (second video) other RNPs. Scale bars: 0.5 µm.

**Video S7. MS2 Reporter RNAs in Live C2C12 Myotubes Imaged for 50 min, Related to Figure 6.** Videos show five myotubes expressing MS2 labeled RNA (top) with RNA trajectories overlaid (bottom). Scale bars: 2 µm.

**Video S8. Computational Simulation Confirms That Directed Transport Is Required to Disperse mRNA in Myofibers, Related to Figure 7.** 10 hr of simulated RNA motion in a segmentation of a real myofiber (see Fig. 1 and S1) in different configurations of diffusive and directed motion states. Videos show 2D projections of three-dimensional simulations. Simulated RNAs in “low-mobility” and “high-mobility” diffusive states are represented as blue and green dots, respectively. “Processive’’ and “Crawling” transport events are shown as magenta and yellow line segments, respectively. Grey dots represent decayed RNAs. Nuclei shown in white. Scale bars: 10 µm.

## METHODS

### RESOURCE AVAILABILITY

#### Lead contact

Further information and requests for resources and reagents should be directed to and will be fulfilled by the Lead Contact, Eric Wang.

#### Materials availability

Plasmids and cell lines generated in this study will be distributed to interested parties upon request.

#### Data and code availability

The code generated during this study is available at https://github.com/cpkelley94/muscle-FISH.

### EXPERIMENTAL MODEL AND SUBJECT DETAILS

#### *Extensor digitorum longus* myofiber isolation

*Extensor digitorum longus* (EDL) muscle was dissected from 10 week old FVB/NJ mice of both sexes. Mouse maintenance and care followed policies advocated by NRC and PHS publications, and approved by Institutional Animal Care and Use Committee (IACUC), University of Florida. Myofiber isolation followed established methods (Pasut et al., 2013). Tissues were digested with prefiltered 0.02% collagenase in DMEM at 37°C for 1 hr. Digested EDLs were flushed by pipetting with pre-warmed DMEM in horse serum (HS)-coated plates under microscopy to dissociate individual fibers. Fibers were collected in HS-coated plates containing DMEM and stored in a 37°C, 5% CO_2_ tissue culture incubator for no more than 1 hr before fixation or continued culture.

#### *Ex vivo* myofiber culture

Isolated myofibers were transferred to HS-coated dishes and cultured in DMEM + 2% HS in a 37°C, 5% CO_2_ tissue culture incubator for up to 36 hr.

#### C2C12 mouse myoblasts

The C2C12 mouse myoblast cell line was obtained from ATCC (CRL-1772). Cells were authenticated via morphological assessment. Myogenic potential was assessed by myotube differentiation followed by morphological assessment (see *C2C12 Myotube differentiation*). Cells were grown in a 37°C, 5% CO_2_ tissue culture incubator on tissue-culture-treated dishes in DMEM + 20% fetal bovine serum (FBS) and passaged with trypsin-EDTA before reaching 70% confluency. Original cells from ATCC were expanded and frozen in DMEM + 20% FBS + 10% DMSO. Cells from frozen stocks were used for no more than 5 passages before being discarded to preserve myogenic potential. For FISH, myoblasts were plated on poly-L-lysine coated coverslips (EMS) at 70% confluency and cultured overnight.

#### Generation of inducible C2C12 myoblast stable cell lines

Stable ponasterone A (PonA)-inducible C2C12 myoblasts were generated using the PiggyBac transposon system by introducing PB-Neo-pERV3 plasmid and mPB PiggyBac transposon plasmid into C2C12 myoblasts at 50% confluency in 6-well plates using Transit-LT1 transfection reagent according to manufacturer’s specifications (Li et al., 2013; Wyborski et al., 2001). Cells were cultured for two days and then treated with 200 µg/mL G418 (Geneticin) antibiotic to select for stable cells.

#### Generation of C2C12 myoblast cell lines for live cell RNA tracking

Stable PonA-inducible C2C12 cells were transduced with lentivirus containing an MS2 coat protein HaloTag fusion protein (MCP-Halo) expression cassette. Cells with MCP-Halo integration were isolated by FACS on a Sony SH800 flow cytometer after staining with JF646 HaloTag ligand (Promega). PB-PuroPonA-Kdm5b-MS2 plasmid and mPB PiggyBac transposon plasmid were then introduced to C2C12 myoblasts at 50% confluency in 6-well plates using Transit-LT1 transfection reagent according to manufacturer’s specifications. Cells were cultured for two days and then treated with 5 µg/mL puromycin antibiotic to select for stable cells.

#### C2C12 myotube differentiation

For differentiation, C2C12 myoblasts were plated at 80% confluency on micromolded gelatin substrates (Bettadapur et al., 2016) prepared on glass bottomed dishes or coverslips for live cell imaging or FISH, respectively. Myoblasts were cultured overnight in DMEM + 20% FBS then switched to DMEM + 2% HS and cultured for an additional 5 days. Media was changed on day 2 and day 4 of differentiation.

### METHOD DETAILS

#### Plasmids and cloning

Plasmid PB-Neo-pERV3 contained bicistronic ecdysone receptor expression cassette from plasmid pERV3 under control of EF1a promoter and PGK-driven neomycin resistance cassette flanked by PiggyBac transposon arms (Li et al., 2013; Wyborski et al., 2001). Plasmid was generated using PCR and Clontech In-Fusion cloning according to manufacturer’s specifications. PB-PuroPonA-Kdm5b-MS2 contained KDM5B coding sequence with 45x MS2 hairpins in the 3’ UTR driven by the ecdysone-responsive promoter (EGSH) and a PGK-driven puromycin resistance cassette. Plasmid was generated using Clontech In-Fusion cloning according to manufacturer’s specifications and standard ligation-based molecular cloning. MS2 hairpin array was derived from plasmid kindly gifted by Edouard Bertrand. Kdm5b coding sequence was derived from plasmid kindly gifted by Tim Stasevich. Plasmid used to generate MCP-Halo lentiviral vectors was a gift from Jeffrey Chao (Addgene #64540).

#### Antibodies

Polyclonal rabbit anti-Fxr1p (13194-1-AP, Proteintech; dilution: 1:200), polyclonal rabbit anti-KIF1C (ab125903, Abcam; dilution: 1:500), polyclonal rabbit anti-Mbnl1 (kindly gifted by Maury Swanson; dilution: 1:1000), monoclonal rabbit anti-Telethonin (ab133646, Abcam; dilution: 1:1000), monoclonal mouse anti-TDP-43 (ab104223, Abcam; dilution: 1:500), monoclonal mouse anti-G3BP (ab56574, Abcam; dilution: 1:1000), monoclonal mouse anti-Alpha Tubulin (T8203, Sigma; dilution: 1:1000), monoclonal mouse anti-Puromycin (EQ0001, Kerafast; dilution: 1:1000), monoclonal mouse anti-Nuclear Pore Complex Proteins (mAb 414, kindly gifted by Maury Swanson; dilution: 1:1000), polyclonal chicken anti-Alpha Tubulin (ab89984, abcam; dilution: 1:1000) were used for protein localization via IF, either alone or in combination with FISH.

#### Drug treatments

Nocodazole was used at 5 μg/mL in myofibers, myoblasts, and myotubes to depolymerize microtubules. Nocodazole washout to allow microtubule re-polymerization was performed in myofibers subsequent to nocodazole treatment. Myofibers were transferred to a new culture dish containing fresh culture medium (DMEM + 2% HS), and washed 3 times with fresh culture medium before continued culture. Partial nocodazole washout to allow formation of a sparse microtubule network was performed subsequent to nocodazole treatment by replacing culture medium with fresh medium within the same culture dish. Actinomycin D was used at 5 μg/mL in myofibers to inhibit transcription. Puromycin was used at 100 μM in myofibers to inhibit translation. Ponasterone A was used at 2 µM in stable C2C12 myotubes to induce MS2 RNA reporter expression.

#### HCR RNA smFISH and immunofluorescence

HCR v3.0 RNA FISH probes for each gene studied were purchased from Molecular Instruments. Primary probe sets contained between 20 and 30 probes, depending on the length of the mRNA. HCR amplifiers and buffers were purchased from Molecular instruments. Freshly isolated myofibers, *ex vivo* cultured myofibers, C2C12 myoblasts, and C2C12 myotubes were processed identically. Samples were fixed in 4% paraformaldehyde (PFA) in phosphate-buffered saline (PBS) for 10 min at room temperature (RT), then washed 3 x 5 min with PBS at RT, followed by permeabilization with 1% Triton X-100 in PBS for 10 min at RT. If immunofluorescence was performed, samples were incubated in PBS containing 1% ultra-pure RNase-free BSA (Sigma), 1 U/µL Nxgen RNase inhibitor (Lucigen), and 0.1% Tween 20 (blocking buffer) for 30 min at RT. Blocking buffer was then replaced with blocking buffer containing diluted primary antibody, and samples were incubated for 1 hr at RT followed by 3 x 5 min washes in PBS + 0.1% Tween 20 (PBST). Samples were then incubated in blocking buffer containing diluted secondary antibodies for 30 min at RT and washed 3 x 5 min in PBST. If FISH was performed, samples were then washed once with PBS and fixed again in 4% PFA for 10 min at RT, followed by 3 x 5 min washes with PBS and 1 x 5 mins wash with 2x saline-sodium citrate (SSC) buffer at RT. Samples were then incubated in pre-warmed HCR hybridization buffer (Molecular Instruments) for 30 min at 37°C in a humidified chamber. During the incubation, primary probes were aliquoted into PCR tubes, heated to 95°C for 90 s, then diluted to 1 nM in pre-warmed hybridization buffer and kept at 37°C until hybridization. Hybridization buffer was replaced with hybridization buffer containing diluted probes, and samples were incubated overnight at 37°C in a humidified chamber. Samples were then washed 5 x 10 min with pre-warmed HCR wash buffer (Molecular Instruments), then washed 2 x 5 min with 5x SSC + 0.1% Tween 20 (SSCT) and incubated in HCR amplification buffer (Molecular Instruments) for 30 min at RT in a humidified chamber. During the incubation, HCR amplifiers were aliquoted into PCR tubes and heated to 95°C for 90 s, then allowed to cool at RT protected from light for at least 30 min. Amplifiers were diluted to 60 nM in HCR amplification buffer just before amplification. Amplification buffer was replaced with amplification buffer containing diluted amplifiers, and samples were incubated for 3 hr at room temperature in a humidified chamber. Samples were then washed 5 x 10 min in SSCT, 1 x 5 min in PBS containing 0.1 µg/mL DAPI, and mounted on slides.

#### Puromycylation

Myofibers were treated with puromycin at 2 μM for 10 min to label nascent peptides. Control myofibers were pre-treated for 30 min with 100 μM anisomycin to inhibit puromycylation. Myofibers were washed 3x with pre-warmed PBS and fixed in 4% PFA in PBS for 10 min.

#### Fixed sample imaging

All myofiber imaging was carried out on a Zeiss LSM 880 AxioObserver microscope with Airyscan using a Plan-Apochromat 1.3 NA 40x oil objective (Fig. 1, 2, 3, 4, 5, S1, S2, S3, S4, S5, S6, Video S2, Video S3), a Plan-Apochromat 1.4 NA 63x oil objective (Fig. S1A, Video S1), or a Plan-Apochromat 1.4 NA 100x oil objective (Fig. S3B and S3C). Airyscan processing was performed on all images in Zeiss ZEN software. Z-stacks were acquired at 0.5 μm intervals. Myoblast and myotube imaging was carried out on a Zeiss LSM 880 AxioObserver microscope using wide field fluorescence illumination, a Plan-Apochromat 1.4 NA 100x oil objective, and Zeiss AxioCam monochrome camera (Fig. 3I). Z-stacks through myoblasts and myotubes were acquired at 0.5 μm intervals.

#### Live cell imaging

Immediately before live imaging, differentiated myotubes were stained with JF646 HaloTag Ligand (Promega) according to manufacturer specifications. Imaging was performed in Fluorobrite phenol-free culture medium at 37°C, 5% CO_2_ in a stage mounted incubator using a Zeiss LSM 880 AxioObserver microscope with Plan-Apochromat 1.46 NA 100x oil objective, widefield fluorescence illumination, and Zeiss AxioCam monochrome camera (Fig. 6). For five fast imaging movies, images were captured continuously for 30 s to 2 min with exposure times between 100 and 500 ms. For long imaging movies, images were captured at 1.0 s exposure time with a 15 s time lag over the course of ∼50 min.

### QUANTIFICATION AND STATISTICAL ANALYSIS

#### General image analysis pipeline

For each FISH experiment, 3D Airyscan confocal stacks in CZI format were first processed using a general automated pipeline to obtain myofiber and nuclei masks and RNA spot coordinates (Fig. S1B). All code is written in Python 3 and is published open source at https://github.com/cpkelley94/muscle-FISH. For each image, voxel dimensions and channel wavelengths were extracted from CZI metadata, and channels were split into 3D arrays. To detect the myofiber geometry, the FISH channel was blurred using a 3D Gaussian kernel, and background signal was binarized using Li’s method for automatic threshold selection (Li and Tam, 1998). Nuclei were detected by thresholding the DAPI channel using a modified Otsu’s method (Otsu, 1979), followed by contraction of the mask by 0.5 μm in each dimension using morphological erosion. Prior to FISH spot detection, cumulative photobleaching along the z-stack in the FISH channel was corrected by fitting an exponential decay model to the average signal intensity along the z-dimension and multiplying the image by the inverse. Spots were detected using the 3D Laplacian of Gaussian method (Lindeberg, 1998), with a kernel scale of 1 voxel and a threshold of 2.5%. To robustly control false positive rate, an automatic signal-to-noise filter was applied, which removed spot calls with a maximum intensity lower than the 90th percentile spot intensity divided by the square root of 10. For each image, fiber segmentation, nuclei detection, and FISH spot detection were inspected by a researcher blind to gene identity, and thresholds were manually adjusted when necessary to maintain accuracy.

For each image, FISH spot density (spots/μm^3^) was calculated by dividing the number of detected spots by the total myofiber volume (Fig. 1D and 1E). The distance from each FISH spot to the nearest nucleus was measured by applying a Euclidean distance transform to the nuclear mask. To generate null distributions for statistical comparison, spot positions were randomized within the cytoplasmic compartment 10,000 times, and the distance to nearest nucleus for each randomized spot was calculated identically as above. The experimental and null distributions were compared using the Mann-Whitney *U* test (Fig. 1F) (Mann and Whitney, 1947). For each gene, the difference in the median distance to nearest nucleus between the experimental and randomized distributions was calculated, and these differences were compared across genes to evaluate correlations with mRNA abundance and length (Fig. S1C)

#### Analysis of FISH signal periodicity

The Gapdh FISH image presented in Fig. 1C was rotated to align the striations with the vertical axis, and the mean FISH signal over the z- and vertical axes was plotted for 40 µm along the fiber. The power spectral density was calculated by discrete fast Fourier transform of the signal from the entire fiber image (Fig. 1G).

#### Analysis of association of RNAs with cytoskeletal filaments

For each gene, image stacks containing FISH of the RNA of interest and IF of filament proteins were first processed using the general pipeline to generate myofiber and nucleus segmentations and FISH spot coordinates. Microtubule and Z-disk segmentations were generated from IF channels (Tuba1a and Tcap, respectively) using the Allen Cell and Structure Segmenter (Chen et al., 2020). Filament segmentations were flattened into 2D by maximum intensity projection along the z-dimension. The 2D distance from each FISH spot to the nearest cytoskeletal filament was measured by applying a Euclidean distance transform to filament segmentations. Spots were considered “cytoskeleton-associated” if they were located within 2 pixels (∼0.1 μm) of either filament mask. To determine if FISH spots were located more proximally to Z-disks and microtubules than expected by chance, a null distribution was generated by randomizing spot coordinates in the cytoplasmic compartment. The experimental and null distributions were compared using the one-sided non-parametric Mann-Whitney *U* test (Fig. 2D).

Z-disk-microtubule intersections (ZMIs) were detected using a novel approach. Flattened Z-disk and microtubule segmentations were skeletonized using Zhang’s method (Zhang and Suen, 1984), and pixels overlapping the nuclear mask were excluded. The two skeletons were merged into a single array with four possible values at each pixel: 0 = background, 1 = Z-disk skeleton, 2 = microtubule skeleton, and 3 = both skeletons. ZMIs were detected by searching the combined mask for a set of small subarrays (motifs) that capture perpendicular intersections. By rotation, reflection, and feature swapping operations, a total of 162 3×3 and 4×4 motifs were algorithmically enumerated from a set of 14 archetypes. Template matching by fast normalized cross-correlation (Briechle and Hanebeck, 2001) was used to efficiently search the combined mask for occurrences of each motif. Multiple intersections called within a radius of 2 px were merged into a single intersection. The distance from each cytoskeleton-associated FISH spot to the nearest ZMI was measured by applying a Euclidean distance transform to the ZMI coordinates. To determine if cytoskeleton-associated FISH spots were located more proximally to cytoskeletal intersections than expected by chance, a null distribution was generated by randomizing spot coordinates in a region of the cytoplasmic compartment within 0.25 μm of either the microtubule or Z-disk skeleton. The experimental and null distributions were compared using the one-sided non-parametric Mann-Whitney *U* test (Fig. 2E).

#### Analysis of RNA spatial patterns after nocodazole treatment

For FISH spots detected in the cytoplasm, the distance to nearest nucleus was calculated as above. For FISH spots detected inside the nucleus, the intranuclear distance was defined as the relative position of the spot between the centroid and periphery of the nucleus. Intranuclear distance was measured by ray casting, using a 3D surface mesh generated from the nuclear mask to approximate the geometry of the nuclear envelope (Fig. 3C, 3D, and S4A). FISH spots were assigned to spatial compartments (nuclear, perinuclear, cytoplasmic) using the aforementioned feature masks, and the fraction of RNAs within the perinuclear compartment was calculated (Fig. 3E). Upregulation of RNAs during culture was assessed by comparing FISH spot density in 18 hr DMSO-treated myofibers to spot density in freshly isolated fibers (Fig. S5C).

#### Estimation of RNA decay rates

For each gene, the mean cytoplasmic FISH spot density was calculated at each time point in the actinomycin D treatment course, and these densities were fit to an exponential decay function using the Levenberg-Marquardt algorithm for nonlinear least squares regression (Fig. S5A and S5B) (Levenberg, 1944). RNA half-life was calculated from the optimized decay constant. The ratio of RNAs >5 μm from the nucleus remaining after 18 hr treatment with nocodazole was calculated and compared to the expected ratio predicted by exponential decay (Fig. 3G).

#### Analysis of perinuclear granule intensity

To enable comparisons of FISH spot intensity across compartments and experimental conditions, spots were first segmented in 3D using the Allen Cell and Structure Segmenter (Chen et al., 2020). Briefly, for each image, the FISH channel was smoothed using a 3D Gaussian kernel with a standard deviation of 1 pixel. Spots were detected using the multi-scale Laplacian of Gaussian method with candidate scales of 1 and 2 pixels, and the transform was thresholded to generate a binary mask. To split merged spots, local peaks in FISH intensity were identified and used as seeds for watershed segmentation within the masked area of the image. For each image, segmented FISH spots were called as “cytoplasmic” if the entirety of the binarized blob overlapped with the cytoplasmic mask, and spots were called as “perinuclear” if the blob at least partially overlapped with the perinuclear mask and did not overlap with the cytoplasmic mask. The intensity of each spot was calculated by integrating the raw FISH signal intensity over all voxels in the binarized blob. Within an image, the relative perinuclear spot intensity was defined as the ratio of the raw spot intensity to the median intensity of cytoplasmic spots (Fig. 4B). The five blobs with the highest intensities were dropped for each RNA/location/condition combination to control for false positive detection, and the 95% confidence interval of the 95th percentile spot intensity was estimated by bootstrapping (Fig. 4C and 4D). Statistical significance at p < 0.05 was determined by inspecting overlap of 95% confidence intervals.

#### Analysis of RNA mobility states

For five short imaging (Fig. 6B and Video S4) and five long imaging (Fig. 6G and Video S7) movies of individual myotubes, time series images were cropped and rotated in FIJI such that the longitudinal axis of each myotube was horizontal. Photobleaching was corrected using histogram matching and background was subtracted using a rolling ball radius of 2 pixels. RNA tracks were obtained using the FIJI TrackMate plugin. Quality and size thresholds for spot detection were set manually using histograms. Linking and gap closing distance were set at 0.5 µm and maximum frame gap was set to 3. Tracks obtained for each movie were inspected and manually linked or unlinked as needed (Fig. 6B and 6G). Trajectories were exported to CSV files and analyzed in Python 3. For each track, spot coordinates were normalized relative to the start position of the track. Then, the maximum distance from the start position to any spot within the track was calculated (Fig. 6H). The percentage of tracks that moved <1 µm (“stationary”) was calculated (Fig. 6I). For five short imaging movies of individual myotubes, tracks were classified into one of four categories based on maximum distance traveled along each axis of the myotube, and the percentage of total tracks in each category was calculated (Fig. 6C and Video S4). Particle trajectories that contained contiguous segments in which a particle moved >1 µm without backtracking were classified as “processive”. The distance from the start and end point of each of these segments was calculated and divided by the elapsed time to obtain the velocity (Fig. 6E). Particle trajectories in which a particle moved at least 1 µm and twice as far in either the X or Y direction were classified as “crawling”. Crawling tracks often moved for only part of the imaging time course and were stationary otherwise. To estimate the velocity of crawling motion, the maximum velocity of any 10 continuous points along crawling tracks was calculated. Maximum distance of the track was used as the distance traveled for “crawling” tracks. Tracks that moved >1 µm and did not travel twice as far in either the X or Y direction were classified as “high mobility” diffusive. Tracks that moved <1 µm were classified as “low mobility” diffusive. For both high and low mobility tracks, mean squared displacement (MSD) was calculated at increasing time lags and a linear least squares regression was performed at the first seven time lags. The slope of the regression line was used to calculate the diffusion coefficient for each particle (Fig. 6D).

#### Markov simulation of RNA transport in myofibers

A discrete-time Markov chain (DTMC) stochastic model was developed in Python and applied to simulate RNA localization dynamics in real myofiber geometries with and without active transport states. Fiber and nuclei masks were segmented from 3D Airyscan microscopy images, and these masks were subtracted to define the cytoplasmic space within which RNAs were allowed to move. During the simulation, RNAs were randomly generated at the periphery of nuclei, with an equal probability of spawning from each nucleus. Once generated, RNAs moved within the cytoplasm according to a set of motion states until degradation (Fig. 7A and Video S8). RNA lifetimes were modeled as exponential particle decay with degradation rate calculated from observed half-life measured using actinomycin D (Fig. S5B). Production of RNAs was modeled as a Poisson process, and production rate was calculated as the product of degradation rate, fiber volume, and mean cytoplasmic spot density observed in FISH images of myofibers. The time-step *t* for the DTMC was 10 s, approximately equal to the average length of directed transport events observed in live-cell imaging.

While moving in the cytoplasm, RNAs randomly transitioned between available motion states, including diffusion (D), crawling transport (C), and processive transport (P). Transition probabilities were constrained by the average number of RNAs observed in each state during live-cell imaging (Fig. 7B). For each RNA, a diffusion coefficient *D* was selected by sampling from the distribution of “low-mobility” diffusion coefficients measured in C2C12 myotubes, smoothed by kernel density estimation (KDE) (Fig. 6D). During each time-step, positions of RNAs in the diffusion state were translated by a 3D Gaussian random variable with a mean of 0 and a standard deviation of (2*Dt*)^1/2^. If allowed, active transport events lasted for a single time-step, and the distance traveled by directed motion (*d_c_* or *d_p_*) was sampled from KDE-smoothed distributions of distances measured in C2C12 myotubes (Fig. 6E). To model the lattice-like structure of microtubules and Z-disks, active transport events were randomly assigned a direction along either the axial or radial dimensions of the fiber, with a 50% probability for each. The axial unit vector was identified by applying a medial axis transform to the fiber mask in 2D and fitting a line to the skeleton by least squares regression. RNAs traveling in the axial direction moved either parallel or antiparallel to the fiber with equal probability. RNAs traveling in the radial direction were allowed to move in any 3D direction perpendicular to the axial unit vector with equal probability. If an RNA exited the allowed cytoplasmic space as a result of its motion during a time-step, the motion event was reverted and sampled again. Particle positions and states were recorded at each time-step for analysis (Fig. 7B).

Using this framework, simulations of Polr2a mRNA transport were conducted for 360,000 time-steps (1000 hr) in the following state configurations: (a) D only, (b) D, C, and P. After discarding the initial 10% of the simulation as burn-in to eliminate non-equilibrium effects, samples were taken every 360 time-steps (1 hr), and the average distribution of RNA across the fiber was computed across all samples (Fig. 7C).

